# Structures of microRNA-precursor apical junctions and loops reveal non-canonical base pairs important for processing

**DOI:** 10.1101/2020.05.05.078014

**Authors:** Grant M. Shoffner, Zhixiang Peng, Feng Guo

## Abstract

Metazoan pri-miRNAs and pre-miRNAs fold into characteristic hairpins that are recognized by the processing machinery. Essential to the recognition of these miR-precursors are their apical junctions where double-stranded stems meet single-stranded hairpin loops. Little is known about how apical junctions and loops fold in three-dimensional space. Here we developed a scaffold-directed crystallography method and determined the structures of eight human miR-precursor apical junctions and loops. Six structures contain non-canonical base pairs stacking on top of the hairpin stem. U-U pair contributes to thermodynamic stability in solution and is highly enriched at human miR-precursor apical junctions. Our systematic mutagenesis shows that U-U is among the most efficiently processed variants. The RNA-binding heme domain of pri-miRNA-processing protein DGCR8 binds longer loops more tightly and non-canonical pairs at the junction appear to modulate loop length. Our study provides structural and biochemical bases for understanding miR-precursors and molecular mechanisms of microRNA maturation.

## Introduction

Canonical microRNAs (miRNAs) in metazoans are transcribed as primary transcripts (pri-miRNAs) ^1, 2^. All pri-miRNAs contain characteristic hairpin secondary structures that contain an imperfectly paired stem and are flanked by unstructured regions ^3, 4^. pri-miRNA hairpin stems consist of approximately 35 ± 1 imperfect base pairs, although substantial variations in stem length occur ^5^. The range of optimal stem length is 33-39 nt when residues in mismatches and internal loops are included in counting ^6^. pri-miRNAs must be recognized and cleaved in the nucleus by the Microprocessor complex that contains the Drosha ribonuclease and its RNA-binding partner protein DGCR8. The cleavage occurs at two staggered sites in a pri-miRNA hairpin stem, producing a precursor miRNA (pre-miRNA) intermediate. The pre-miRNA contains the upper ∼2/3 of the pri-miRNA hairpin with 2- or 1-nt 3’ overhang. The pre-miRNA is then exported to the cytoplasm and cleaved by another ribonuclease Dicer at two staggered sites close to the apical loop, resulting in a miRNA duplex. One or both strands of the duplex can be incorporated into the effector Argonaute proteins and become mature miRNAs functional in regulating gene expression ^1^. Pri- and pre-miRNAs are collectively called miRNA precursors or miR-precursors in short.

The hairpin loops of miR-precursors, most often called apical loops but sometimes terminal loops, are important features for processing. Apical loops are preferred to be longer than 9 nt for pri-miRNAs to be efficiently cleaved by Microprocessor ^7^. A more recent genomic study suggested that the optimal loop length is 8-16 nt ^6^. Mutagenesis of the loop and natural genetic variation in this region can have severe impacts on processing efficiency and cleavage site selection ^8, 9^. The junction between miRNA hairpin stem and apical loop, called the apical junction, serves as an anchor point for Microprocessor to determine the cleavage sites ^5, 10^, along with the basal junction at the other end of pri-miRNA hairpin stem ^4^. The apical junctions and loops have been shown to be important for pre-miRNA cleavage by Dicer ^8, 11^. Furthermore, a growing list of regulatory proteins modulate miRNA processing at both Drosha and Dicer cleavage steps by association with apical loops ^12, 13, 14^.

The recognition of pri-miRNAs by Microprocessor serves a gate-keeping function for the miRNA maturation pathway. Microprocessor is a heterotrimeric complex containing one Drosha polypeptide chain and a DGCR8 dimer ^15, 16^. It clamps the pri-miRNA hairpins from both ends ^3, 4, 7, 10^, with DGCR8 binding in the apical region and Drosha in the basal region ^15^. DGCR8 contains a dimeric RNA-binding heme domain (Rhed) that specifically binds the apical junction ^17^. Among known RNA-binding domains, Rhed is unique in that it requires a heme cofactor with the Fe center in the 3^+^ state ^18, 19, 20, 21, 22^. A UGU motif at the 5’ end of the apical loop has been shown to enhance processing ^23^. The heme cofactor is important for proper orientation of pri-miRNA hairpins in Microprocessor and for preferentially processing of distinct groups of pri-miRNAs, including those containing the UGU motif ^24, 25, 26^. Most recently, cryo-EM structures of Microprocessor in complex with pri-miRNAs have revealed an extensive protein-RNA interface, but pri-miRNA apical junctions and loops and the Rhed domain could not be resolved ^27, 28^.

Our understanding of miRNA hairpins is mostly based on secondary structure predictions ^29^, sometimes assisted by chemical or nuclease footprinting ^30^. We are interested to determine three-dimensional structures of apical junctions and loops for their important roles in miRNA maturation and regulation ^5, 7, 8, 10^ and as targets for drug discovery ^31^. Apical junctions and loops are shared between pri-miRNA and pre-miRNA, but here for simplicity we use pri-miRNA to name individual apical junctions and loops. To date only two apical stem-loops have been structurally characterized, using NMR spectroscopy, with pri-miR-20b folding to well-defined rigid structures ^32^ and pri-miR-21 unstructured ^31, 33^. The human genome encodes 1,881 pri-miRNA hairpins that differ from each other greatly in terms of both their sequences and predicted secondary structures ^29^. Toward surveying the large number of miR-precursor structures, we develop a new technique that enables rapid determination of hairpin loop structures and report nine apical junction and loop structures from eight pri-miRNAs. We interpreted these structures with help from sequence analysis, biochemical, and cellular assays and revealed previously unknown principles governing miRNA maturation.

## Results

### Survey of miR-precursor apical loop length

A previous investigation showed that pri-miRNAs with short (<10 nt) apical loops tend to be processed inefficiently by Microprocessor ^7^. We compiled a list of human pri-miRNA apical loop sequences based on predicted secondary structures we produced using mfold ^34^ and similar ones provided by miRBase ^29^. Majority of them (1,314 out of 1,881, 70%) are less than 10 nt long, with the highest frequencies in the 4-6 nt range (Fig. 1a). RNA secondary structure prediction programs tend to include base pairs in relatively long loops that are not necessarily stable ^31, 32^. We disregarded 1 or 2 base pairs that are separated from the hairpin stem on both strands. Although the list might still underestimate the number of longer loops, it nevertheless reflects our best knowledge. Therefore, for most miR-precursor recognition events, the processing machinery such as the Rhed in DGCR8 and the helicase domain in Dicer must interact with a relatively short apical loop in order to access the apical junction.

**Fig. 1.**
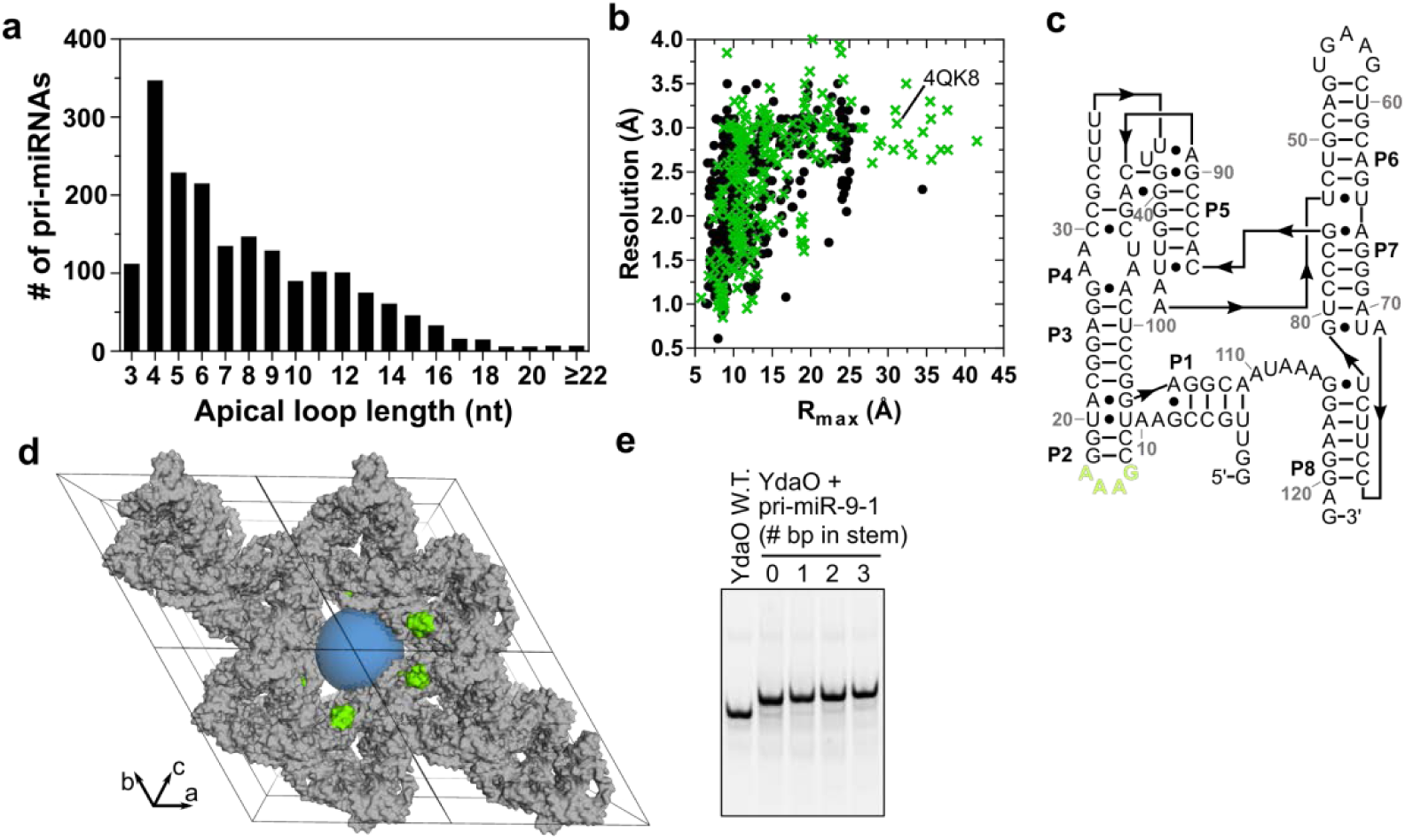
Analysis of pri-miRNA terminal loops and search for potential crystallization scaffolds. **a** Distribution of pri-miRNA apical loop lengths. **b** Comparison of the largest spherical cavity (with radius R_max_) present in each RNA crystal structure against the diffraction resolution of the structure. Crystal forms with a single chain in the asymmetric unit are shown as green crosses, and all others as black dots. **c** Secondary structure of the YdaO-type ci-di-AMP riboswitch. **d** Crystal packing of the riboswitch (PDB ID 4QK8). Molecules surrounding a large central channel (parallel to the *c*-axis) are colored gray, and a blue sphere with a radius of 31 Å is positioned in the channel to illustrate its size. The L2 stem loops terminating inside the channel are green. **e** Native gel analysis of WT YdaO and fusions with the pri-miR-9-1 terminal loop with 0-3 base pairs from the stem.

### Scaffold-directed crystallography

To determine the three-dimensional structures of pri-miRNA apical junctions and loops, we developed a scaffold-directed crystallization approach. The concept is to fuse the target (unknown) sequence onto a scaffold molecule known to crystallize well and with a crystal structure available. The fusion should crystallize under conditions similar to that for the scaffold alone. The crystal lattice should be able to accommodate the target moiety. The scaffold structure allows the structure of the fusion to be determined via molecular replacement.

To identify a suitable scaffold, we mined the Protein Data Bank (PDB) for RNA crystals fulfilling four criteria. For each RNA structure entry, we first identified the largest sphere that can be accommodated in the lattice cavity, as characterized by the radius *R*_max_ (Fig. 1b). We considered the diffraction resolution reported. To simplify the design, we focused the search on entries with one RNA chain in the asymmetric unit, but also considered select candidates with multiple chains. Finally, we manually reviewed the crystal lattices to find stem-loops that point toward the lattice cavity so that an RNA hairpin can be fused to. Amongst hundreds of structures surveyed, we identified only one RNA meeting these requirements, the YdaO-type c-di-AMP riboswitch from *Thermoanaerobacter pseudethanolicus* (abbreviated from here on as YdaO) ^35^.

The YdaO crystal lattice contains large solvent channels with *R*_max_ ∼30 Å. The riboswitch has a complex pseudo-two-fold symmetric ‘cloverleaf’ fold (Fig. 1c). Its short P2 stem is positioned inside the channel and substantially away from neighboring molecules (Fig. 1d). P2 is capped by a GAAA tetraloop, which has strong and concrete electron density for every residue. The structure of this tetraloop is very similar to those determined in previously studies, indicating little interference of its folding by the scaffold, crystal lattice, crystallization or cryoprotection conditions. We replaced the tetraloop with the 14-nt pri-miR-9-1 apical loop plus 0-3 additional base-pairs from the stem. All four fusion RNAs, after annealing in the presence of the c-di-AMP ligand, migrated as single bands on a native gel (Fig. 1e), indicating that the engineered pri-mi-RNA sequences do not interfere with the scaffold folding. These properties make P2 an excellent site for grafting hairpins.

For our representative set of short pri-miRNA loops, we generated fusions with the YdaO scaffold containing the loop plus a various number of base pairs from the stem, and screened for crystallization. We succeeded in obtaining crystals for constructs containing 1 or 0 base pair from the pri-miRNA stems, annotated as “+1bp” and “+0bp” respectively. These crystals belong to the same space group, P3_1_21, with similar cell dimensions (Table 1). We collected X-ray diffraction data and determined nine structures at resolutions ranging from 2.71 to 3.10 Å. The resolutions of seven structures are higher than the previously reported 3.05 Å of the wild-type (WT) YdaO ^35^. For four pri-miRNAs, we also collected single-wavelength anomalous dispersion (SAD) data with redundancy in the 79-115 range. These SAD data improved the quality of electron density maps and were included in refinement. The refined native structures showed that the scaffold moieties are very similar to the WT, with C1’ root-mean-square deviation (RMSD) values ranging from 0.22 to 1.18 Å. Below we describe the pri-miRNA moieties. Unlike many RNA loop structures in PDB, our structures are free from crystal contacts or interactions with ligands, and thereby reflect their own folding propensities.

**Table 1.** Data collection and refinement statistics for pri-miRNA loop fusion structures.

| RNA | 378a+0bp |  | 378a+1bp | 340+1bp | 300+0bp | 202+1bp |  | 208a+1bp | 449c+1bp |  | 320b-2+1bp |  | 19b-2+1bp |
| --- | --- | --- | --- | --- | --- | --- | --- | --- | --- | --- | --- | --- | --- |
| PDB code | 6N5T |  | 6N5Q | 6N5P | 6WTR | 6N5O |  | 6N5N | 6N5K |  | 6N5S |  | 6WTL |
| Data Collection |  |  |  |  |  |  |  |  |  |  |  |  |  |
| Data set | Native | SAD | Native | Native | Native | Native | SAD | Native | Native | SAD | Native | SAD | Native |
| No. of crystals | 1 | 1 | 1 | 1 | 1 | 2 | 1 | 1 | 1 | 2 | 1 | 3 | 1 |
| Wavelength (Å) | 1.116 | 1.907 | 1.116 | 1.116 | 1.116 | 0.9202 | 1.907 | 0.9791 | 0.9791 | 1.907 | 0.9202 | 1.907 | 0.9791 |
| Data range (°) | 360 | 3,240 | 360 | 720 | 360 | 1,080 | 4,320 | 180 | 360 | 4,320 | 720 | 4,920 | 360 |
| Space group | P3<sub>1</sub>21 | P3<sub>1</sub>21 | P3<sub>1</sub>21 | P3<sub>1</sub>21 | P3<sub>1</sub>21 | P3<sub>1</sub>21 | P3<sub>1</sub>21 | P3<sub>1</sub>21 | P3<sub>1</sub>21 | P3<sub>1</sub>21 | P3<sub>1</sub>21 | P3<sub>1</sub>21 | P3<sub>1</sub>21 |
| a,b,c (Å) | 113.8,<br>113.8,<br>115.0 | 114.0,<br>114.0,<br>115.1 | 114.9,<br>114.9,<br>114.7 | 114.8,<br>114.8,<br>115.6 | 113.1,<br>113.1,<br>114.1 | 114.9,<br>114.9,<br>115.3 | 115.1,<br>115.1,<br>115.3 | 114.7,<br>114.7,<br>114.8 | 114.0,<br>114.0,<br>114.8 | 115.0,<br>115.0,<br>115.1 | 114.6,<br>114.6,<br>115.1 | 114.9,<br>114.9,<br>115.6 | 115.3, 115.3,<br>115.3 |
| α, β, γ (°) | 90, 90,<br>120 | 90, 90,<br>120 | 90, 90, 120 | 90, 90,<br>120 | 90, 90,<br>120 | 90, 90,<br>120 | 90, 90,<br>120 | 90, 90, 120 | 90, 90,<br>120 | 90, 90,<br>120 | 90, 90,<br>120 | 90, 90,<br>120 | 90, 90, 120 |
| Resolution (Å) | 74.8 –<br>2.79<br>(2.89 –<br>2.79) | 74.9 –<br>3.18<br>(3.29 –<br>3.18) | 75.1 – 2.95<br>(3.05 –<br>2.95) | 75.4 –<br>2.99<br>(3.10 –<br>2.99) | 74.3 –<br>3.08<br>(3.19 –<br>3.08) | 99.5 –<br>2.71<br>(2.81 –<br>2.71) | 75.4 –<br>2.95<br>(3.05 –<br>2.95) | 75.1 – 2.95<br>(3.03 –<br>2.95) | 74.9 –<br>3.10<br>(3.18 –<br>3.10) | 75.3 –<br>3.59<br>(3.71 –<br>3.59) | 75.1 –<br>2.80<br>(2.87 –<br>2.80) | 99.5 –<br>3.17<br>(3.28 –<br>3.17) | 75.5 – 2.85<br>(2.92 – 2.85) |
| R<sub>meas</sub>(%)<sup>†</sup> | 5.9<br>(140) | 16.7<br>(195) | 6.7<br>(154) | 10.1<br>(202) | 9.2<br>(143) | 10.6<br>(217) | 9.6<br>(146) | 9.0<br>(245) | 11.2<br>(233) | 10.6<br>(189) | 6.2<br>(173) | 16.0<br>(170) | 6.1 (162) |
| R<sub>p.i.m.</sub>(%)<sup>†</sup> | 1.4<br>(32.7) | 1.4<br>(16.0) | 1.5 (34.6) | 1.6<br>(32.4) | 2.1 (34.4) | 1.4<br>(33.6) | 0.7<br>(17.0) | 2.0 (47.6) | 1.9<br>(44.4) | 0.8<br>(22.2) | 2.0<br>(49.9) | 1.1<br>(20.9) | 1.4 (35.8) |
| <I/σ> | 27.7<br>(2.1) | 31.4<br>(3.9) | 25.5 (1.9) | 27.1<br>(2.4) | 18.9 (1.7) | 29.9<br>(2.1) | 64.2<br>(3.5) | 24.9<br>(1.9) | 26.6<br>(1.8) | 44.8<br>(2.5) | 23.0<br>(1.4) | 42.1<br>(2.9) | 32.8 (2.0) |
| CC<sub>1/2</sub> (%) | 100<br>(84.8) | 100<br>(87.6) | 100 (84.0) | 100<br>(84.8) | 100 (90.3) | 99.6<br>(81.5) | 100<br>(93.3) | 99.9 (83.0) | 100<br>(86.6) | 100<br>(95.0) | 99.8<br>(70.9) | 99.9<br>(92.4) | 99.9 (81.6) |
| Completeness (%) | 99.1<br>(88.5) | 99.9<br>(99.1) | 99.8 (97.5) | 99.9<br>(100) | 99.2 (92.4) | 99.9<br>(99.0) | 99.9<br>(99.7) | 99.9 (99.0) | 100<br>(100) | 99.5<br>(95.9) | 99.9<br>(99.7) | 99.7<br>(96.7) | 100.0 (99.8) |
| Redundancy | 20.0<br>(17.0) | 79.1<br>(67.8) | 19.4 (18.9) | 38.2<br>(35.7) | 18.8 (15.3) | 56.4<br>(39.1) | 111<br>(25.7) | 19.9 (20.7) | 39.8<br>(39.2) | 100<br>(34.2) | 9.92<br>(9.95) | 115<br>(29.9) | 19.8 (17.9) |
| No. of unique reflections | 21,687 | 14,907 | 18,836 | 18,192 | 15,789 | 24,343 | 19,022 | 18,748 | 16,089 | 19,867 | 21,893 | 28,896 | 21,107 |
| Refinement |  |  |  |  |  |  |  |  |  |  |  |  |  |
| Resolution (Å) | 74.8 – 2.79 |  | 75.1 – 2.95 | 75.4 –<br>2.99 | 74.3 –<br>3.08 | 99.5 – 2.71 |  | 75.1 – 2.95 | 74.8 – 3.10 |  | 75.1 – 2.80 |  | 75.5 – 2.85 |
| R<sub>work</sub> / R<sub>free</sub> | 0.180 / 0.196 |  | 0.165 /<br>0.188 | 0.151 /<br>0.181 | 0.165 /<br>0.192 | 0.180 / 0.219 |  | 0.168 /<br>0.186 | 0.177 / 0.186 |  | 0.177 / 0.201 |  | 0.187 / 0.213 |
| No. Atoms |  |  |  |  |  |  |  |  |  |  |  |  |  |
| RNA | 2,705 |  | 2,747 | 2,722 | 2,676 | 2,705 |  | 2,682 | 2,681 |  | 2,663 |  | 2,661 |
| Ligand | 88 |  | 88 | 88 | 88 | 88 |  | 88 | 88 |  | 88 |  | 88 |
| Ions | 10 |  | 10 | 10 | 9 | 10 |  | 14 | 8 |  | 11 |  | 10 |
| Water | 0 |  | 4 | 4 | 4 | 4 |  | 0 | 6 |  | 6 |  | 1 |

<sup>1</sup>R-factors are defined as:

### Structures of miR-precursor apical junctions and loops

The nine structures cover eight pri-miRNA sequences. pri-miR-378a has two structures, with 0 and 1 canonical base pair from the hairpin stem respectively. Six others contain 1 base pair from the stem and one (pri-miR-300+0bp, or simply 300+0bp) has none. The last base pair of pri-miR-300 stem is identical to the terminal C-G pair of the scaffold, thus the 300+0bp structure is effectively that of 300+1bp. Therefore, our structures reveal minimal apical junctions for all eight pri-miRNAs.

Our series of pri-miRNA structures cover the most frequent loop lengths in humans, ranging from 4 to 8 nt. The longest loop is 8 nt, from pri-miR-378a. As RNA loops can be flexible, they are often not well resolved in the electron density. To our surprise, the electron density maps of both 378a+0bp and 378a+1bp reveal highly structured conformations with clear density for all residues (Fig. 2a,c and Supplementary Fig. 1a,b). The two independently determined loop structures are in close agreement with each other, with RMSD of 1.4 Å over all non-hydrogen atoms. Both structures clearly show that outermost residues in the loop sequence, C1 and A8, form a non-canonical pair mediated by a single hydrogen bond between C1^O2^ and A8^N6^ (Fig. 2b,d). The C-A pair creates a platform onto which remaining loop bases stack. On the 5’ end, two pyrimidine residues C2 and U3 stack above C1. From the 3’ side, four layers of purine residues A4, G5, A6, and A7 stack above A8. The turn bridging the pyrimidine and purine stacks is stabilized by a hydrogen bond between U3^O2’^ and G5^N7^ (Fig. 2b,d). C2^O2^ and A7^N6^ are 3.2 Å apart in the 378a+1bp structure and 3.6 Å in 378a+0bp, potentially forming an additional hydrogen bond across the two base stacks and stabilizing the loop. Interestingly, the 378a loop stacks on a C-G pair in 378a+0bp and the natural A-U pair in 378a+1bp. The fact that these structures are very similar suggests that the pri-miR-378a loop conformation is not strongly influenced by the terminal base pair from the hairpin stem.

**Fig. 2.**
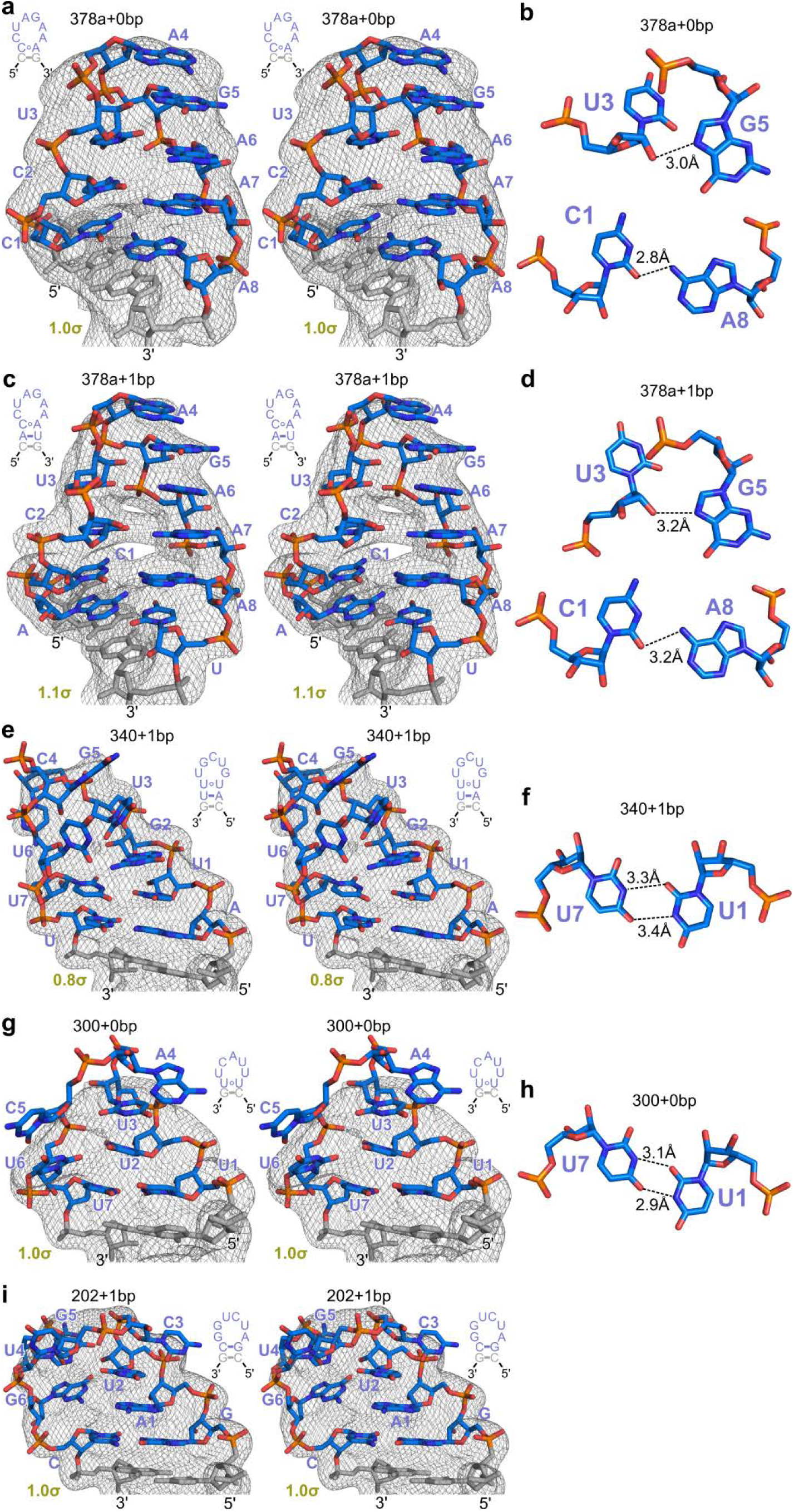
Atomic structures of pri-miRNA apical junctions and loops 8-6 nt in length determined by scaffold-directed crystallography. Throughout the Figure, the last C-G pair from the scaffold P2 stem is colored gray. Panels **a, c, e, g**, and **i** show the labeled structures in stereographic view with σ_A_-weighted 2Fo-Fc electron density maps contoured at levels indicated in each panel. Insets show the sequences and secondary structures. **a** pri-miR-378a (378a+0bp). **b, d, f, h** Hydrogen bonds in the 378a+0bp, 378a+1bp, 340+1bp, and 300+0bp loop structures. The neighboring canonical pair in pri-miR-300 is C-G, identical to the pair in the scaffold. Therefore, the structure is essentially 300+1bp.

The structures of pri-miR-340 (340+1bp) and pri-miR-300 (300+0bp) contain 7-nt loops. The 340+1bp structure confirms the presence of the terminal A-U pair, which is capped by a previously unexpected U1-U7 pair (Fig. 2e,f). These residues are indicated by strong electron density in simulated annealing omit map (Supplementary Fig. 1c). The density gradually grows weaker as the loop extends further away from the stem. The G2 base from the 5’ end of the loop stacks on top of the U-U pair. At the top of the loop, U3, C4, G5, and U6 occupy partial density and are likely in a more flexible conformation. The 300+0bp structure shows non-canonical pairing between U1 and U7, in a geometry similar to that in 340+1bp (Fig. 2g,h and Supplementary Fig. 1d). U2 and U3 stack on U7 across the loop, instead of the neighboring U1 like in the pyrimidine stack in pri-miR-378a, nevertheless ordering the 5’ half of the loop. U6 does not form another non-canonical pair with U2, probably due to backbone constraints. A4 and C5 are to a large extent outside the density and appear to be more flexible.

In the structure of pri-miR-202 (6-nt loop), we did not observe non-canonical base pairs. However, similar to other structures, the A1 and U2 bases at the 5’ end of the loop stack to the final G-C pair of the pri-miR-202 stem (Fig. 2i and Supplementary Fig. 1e). The rest of the loop shows continuous electron density at 1σ, but we could not determine the conformation with high confidence. Overall, the structures of the relatively long (6-8-nt) pri-miRNA loops reveal extensive base stacking and non-canonical base pairing interactions, perhaps stabilizing the loops more than previously anticipated. As a consequence, fewer loop residues are conformationally flexible.

Next, we investigated the structures of shorter pri-miRNA terminal loops (4-5 nt, Fig. 3). The structure of pri-miR-208a (208a+1bp) with a 5-nt loop revealed an unpredicted A1-U5 Hoogsteen pair positioned above the final G-C pair from the stem (Fig. 3a,b and Supplementary Fig. 2a). U2 stacks with a tilt onto the A1 base in the Hoogsteen pair. C5 and G4 possibly further stack on U2, but these residues are partially out of density. In the structure of pri-miR-449c (449c+1bp), U1 and U5 recapitulate the non-canonical U-U pairs from pri-miR-340 and pri-miR-300 (Fig. 3c,d and Supplementary Fig. 2b). G2 stack above U1. The backbones of A3 and U4 are defined by continuous density, but we are less confident about the positions of their bases. The two pentaloops share a theme: the first and fifth residues form non-canonical base pairs, whereas one or more of the remaining, unpaired residues stack on the 5’ side.

**Fig. 3.**
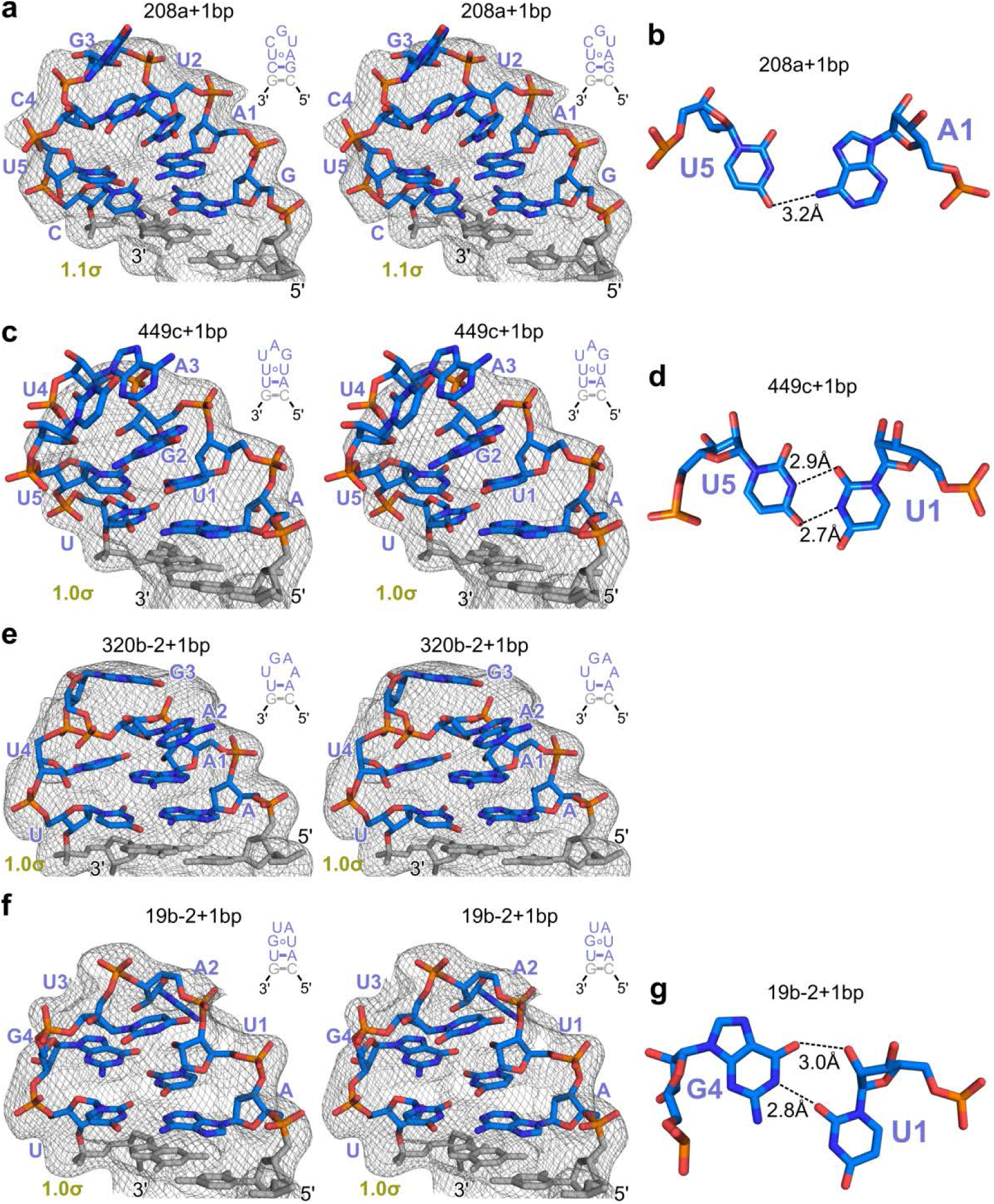
Structures of pri-miRNA apical junctions and loops 4-5 nt in length. Color scheme is identical to that in Fig. 2. **a, c, e, f** Structures and σ_A_-weighted 2Fo-Fc maps of pri-miR-208a (208a+1bp), pri-miR-320b-2 (320b-2+1bp), pri-miR-449c (449c+1bp), and pri-miR-19b-2 (19b-2+1bp). **b, d, g** Non-canonical pairs in the 208a+1bp, 449c+1bp, and 19b-2+1bp loop structures.

In the AAGU tetraloop of pri-miR-320b-2 (320b-2+1bp), we observed continuous electron density for the backbone and the 5’ A1 base stacking atop the terminal A-U pair of the stem (Fig. 3e, Supplementary 2c). The positions of the other three bases are not well-defined. The AAGU loop is also found in non-canonical substrates of yeast RNase III Rnt1 and its structures with and without Rnt1 have been solved using NMR ^36, 37^. Our structure is similar to the structures of Rnt1 substrate despite the different terminal stem base pair (C-G) in the latter.

Finally, the electron density maps of 19b-2+1bp are consistent with its UAUG tetraloop adopting a Z-turn conformation typically seen in the highly stable UNCG tetraloops (N represents any residues) ^38^ (Fig. 3f and Supplementary Fig. 2d). U1 and G4 form a non-canonical pair that involves two hydrogen bonds, U1^O2^—G4^N1^ and U1^O2’^—G4^O6^ (Fig. 3g), with the G4 residue in a *syn* conformation. The U1-G4 pair stacks to the terminal A-U pair in the stem with mostly U1 overlapping. This stacking is less optimal and thereby likely to be less stable than that in the structure of UNCG stacking on a terminal G-C pair, as it is previously shown that a UNCG loop stacking on a G-C pair is more stable than on a C-G pair by 2.4 kcal/mol ^39^. The U3 base stacks on top of U1, with its ribose in C2’-endo sugar pucker. Overall, the 4-5-nt loop structures confirm that the non-canonical pairing and base stacking of the 5’ loop residues witnessed in longer loop structures also dominate the folding of the shorter loops.

### A structural consensus of miR-precursor apical junctions

Our miR-precursor stem-loop structures point toward a set of structural features common to most apical junctions. We further illustrate these features in a structural alignment of all eight miR-precursors (378a+1bp was used to represent pri-miR-378a) (Fig. 4a). First, we always observe the mfold-predicted canonical base pair at the apical end of pri-miRNA stem (5’-0 paired with 3’-0). Because the loops are of different sizes, here we use 5’-0 as the starting point for counting residues from 5’-end of miR-precursor loop sequence, and 3’-0 for counting residues from 3’-end. Second, in all structures the first nucleotide on the 5’ end of the loop base-stacks with the terminal base pair (5’-1 stacking with 5’-0/3’-0). Third, in six of the eight loops (378a, 340, 300, 208a, 449c, and 19b-2), this base stacking is also accompanied by a non-canonical base pair (5’-1 pairing with 3’-1), effectively shortening the apical loops by two nucleotides. Fourth, all eight structures reveal at least one additional level of base-stacking interactions on the 5’ side. For seven of them, it is 5’-2 stacking on 5’-1. For the 19b-2 tetraloop, 5’-3 (U3) stacks on 5’-1, whereas the 5’-2 (A2) base points away. On the 3’ side, only one miR-precursor 378a has second-layer stacking. Beyond these common features, other residues of the miR-precursor loops appear to adopt quite different conformations or are flexible.

**Fig. 4.**
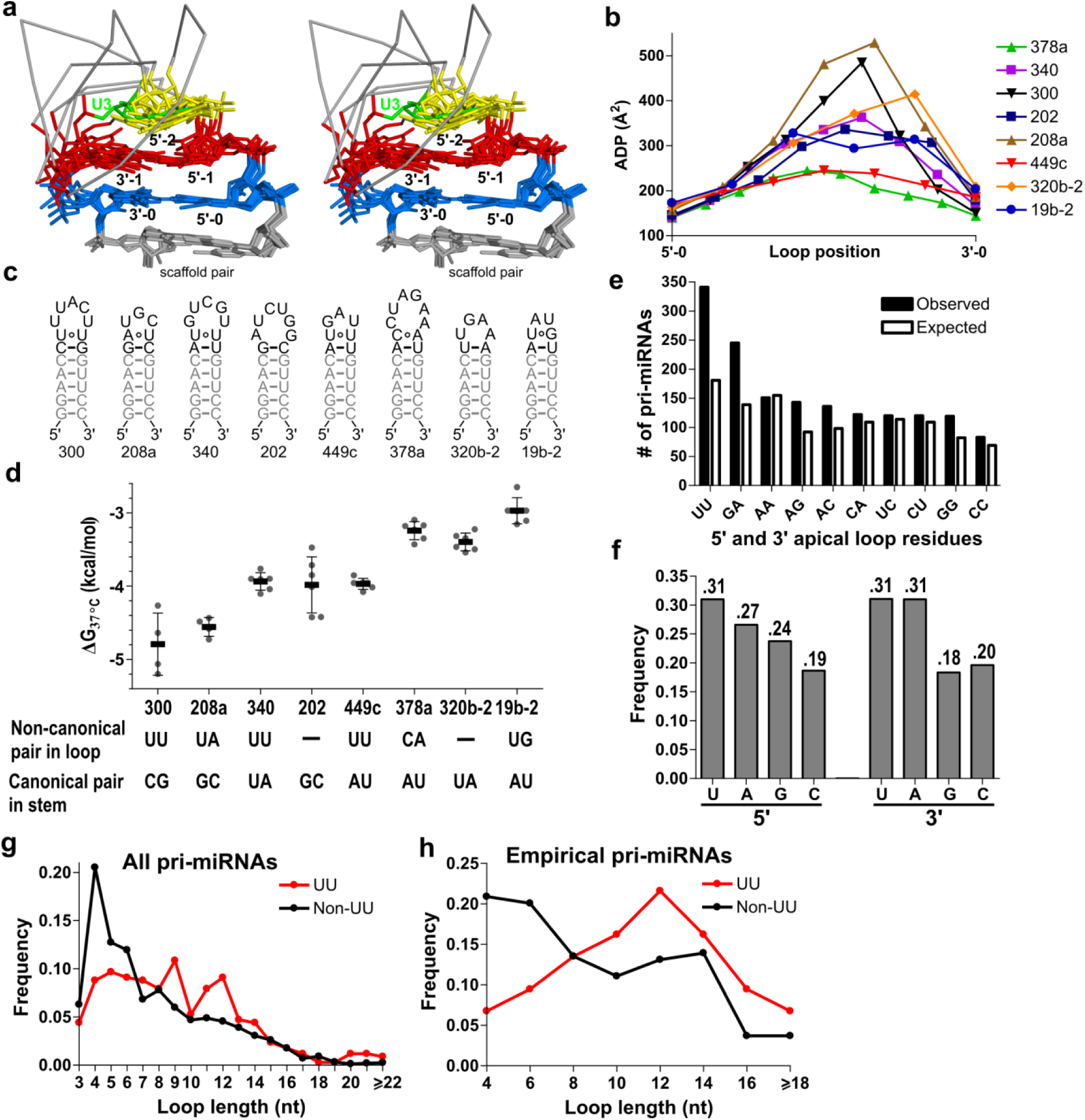
Structural consensus, non-canonical pairs, and asymmetric flexibility of human pri-miRNA apical junctions and loops. **a** Structural alignment of all eight loops. Positions that align well among most or all of the structures are labeled. **b** Average atomic displacement parameter (ADP) per residue, with all loops plotted on the same scale. The 5’ and 3’ ends represent the terminal base pair of the pri-miRNA stem loop. Structure drawings illustrating ADP distribution are presented in Supplementary Fig. 3. **c** Secondary structures of RNA constructs used for optical melting assays. Each RNA contains the pri-miRNA apical loop and the immediately neighboring base pair from the stem, along with five common base pairs shown in gray. **d** Plot of folding ΔG values (mean ± SD) for the eight pri-miRNA apical junctions and loops measured with 50 mM NaCl. Experiments for all RNAs were repeated 6 times except 300 and 208a, which had 4 replicates each. Individual data points are shown as gray dots. Detailed thermodynamic parameters and their standard deviations are listed in Table 2. **e** Observed and expected counts of human pri-miRNAs with the indicated apical loop-closing residue pairs. The expected counts are estimated based on the frequencies of 5’ and 3’ loop residues as shown in **f. g** Loop length distribution of pri-miRNAs with apical loops closed by UU vs non-UU residues. Frequency is defined as fraction of total. **h** Loop length distribution for the set of empirical pri-miRNAs ^6^. To reduce noise, loop lengths are binned. For example, “4” on the graph represents loop lengths of 3 and 4.

### Asymmetric stability of miR-precursor apical loops

Structural stability and dynamics are likely to be important for miR-precursor junctions and loops for at least two reasons. First, common conformational features are expected to be stable. Second, dynamic regions make it easier to avoid steric hindrance when binding processing proteins and to adopt conformations favorable for processing. To investigate this, we first reviewed the atomic displacement parameters (ADPs, also known as the temperature factors or B-factors) refined during structure determination. Not surprisingly, residues at the top of the loop have large ADPs, suggesting that they are highly dynamic; whereas residues close to the stems, which are often involved in common structural features such as non-canonical pairs and base stacking, tend to have lower ADPs (Supplementary Fig. 3). Importantly, most loops display higher stability in their 5’ regions, likely due to stronger stacking interactions. The only exception is pri-miR-378a, in which the 3’-purine stack is slightly more stable than the 5’ pyrimidine stack. To further compare ADPs between structures, we calculated the average ADP for each residue and then plotted the values along the loop positions (Fig. 4b). The peak in ADPs is consistently located in the 3’ half of the loop across most structures. The asymmetric stability of miR-precursor loops is further supported by the generally stronger electron densities for residues at the 5’ end (Fig. 2 and 3, and Supplementary Fig. 1 and 2). It is interesting to note that the UGU motif previously identified to promote efficient processing ^5, 23, 25^ is located at the more stable 5’ end of the loop.

**Table 2.** Thermodynamic parameters for pri-miRNA apical junction and loop folding at 50 mM NaCl, reported as ± standard deviation (n = 4-6).

| RNA<br>(terminal bp) | Non-canonical<br>pair | Loop<br>length<br>(nt) | $T_M$<br>(°C) | $\Delta H$<br>(kcal/mol) | $\Delta S$<br>(e.u.) | $\Delta G_{37^\circ C}$<br>(kcal/mol) |
| --- | --- | --- | --- | --- | --- | --- |
| pri-miR-378a<br>(A-U) | C-A | 8 | 62.9 ± 0.8 | -42.1 ± 1.6 | -125.4 ± 4.9 | -3.24 ± 0.12 |
| pri-miR-340<br>(U-A) | U-U | 7 | 64.9 ± 0.5 | -47.7 ± 1.3 | -141.1 ± 3.7 | -3.94 ± 0.12 |
| pri-miR-300<br>(C-G) | U-U | 7 | 72.4 ± 0.6 | -46.8 ± 4.6 | -135.5 ±<br>13.4 | -4.79 ± 0.42 |
| pri-miR-202<br>(G-C) | — | 6 | 69.5 ± 0.3 | -42.1 ± 4.3 | -122.8 ±<br>12.6 | -3.98 ± 0.38 |
| pri-miR-208a<br>(G-C) | U-A | 5 | 72.2 ± 0.5 | -44.7 ± 1.5 | -129.5 ± 4.4 | -4.56 ± 0.13 |
| pri-miR-449c<br>(A-U) | U-U | 5 | 70.8 ± 0.5 | -40.4 ± 1.0 | -117.5 ± 2.9 | -3.97 ± 0.08 |
| pri-miR-320b-2<br>(U-A) | — | 4 | 72.3 ± 0.7 | -33.3 ± 1.4 | -96.4 ± 4.1 | -3.40 ± 0.12 |
| pri-miR-19b-2<br>(A-U) | U-G | 4 | 70.6 ± 1.0 | -30.4 ± 1.2 | -88.5 ± 3.2 | -2.97 ± 0.18 |

### Non-canonical base pairs contribute to thermodynamic stability

We next asked if the structures of apical junctions and loops we observed contribute to their stability in solution. This question is particularly important as the crystals were frozen in a cryoprotectant solution containing relatively high concentrations of salts (0.2 M (NH_4_)_2_SO_4_, 0.2 M Li_2_SO_4_ in 0.1 M NaHEPES buffer). The salt concentrations in crystallization conditions were even higher (1.63 – 1.90 M (NH_4_)_2_SO_4_, 0.2 M Li_2_SO_4_ in 0.1 M NaHEPES buffer) and may also influence the RNA structures. To investigate this question, we fused the eight pri-miRNA sequences to a common 5-bp helical segment (Fig. 4c) and measured their thermodynamic parameters using optical melting. Like in the crystal structure, each pri-miRNA sequence contains the apical loop and an immediately neighboring canonical base pair from the stem so that a minimal apical junction is included. We expect that the canonical stem base pairs contribute differentially to the overall stability, with the G-C or C-G pairs in three pri-miRNAs being more stable than the A-U and U-A pairs in the others. However, this difference does not fully explain the free energy changes (ΔG) of folding we measured (Table 2). When we take into account the non-canonical pairs we revealed in the three-dimensional structures, a trend appears. The two pri-miRNAs that have G-C or C-G as terminal stem pairs and form non-canonical pairs (pri-miR-300 and pri-miR-208a) are the most stable, whereas the ones that contain A-U or U-A canonical stem pairs but do not form non-canonical base pairs or very weak ones defined by a single hydrogen bond (pri-miR-320b-2 and pri-miR-378a) are among the least stable (Fig. 4d). Most other pri-miRNA sequences that either contain non-canonical pairs but A-U/U-A stem pairs (pri-miR-340 and pri-miR-449c), or form no non-canonical pairs but with G-C/C-G stem pairs (pri-miR-202) are intermediate in stability. The pri-miR-19b-2 apical junction/loop is least stable despite the presence of a U-G pair in the loop but as mentioned above this may be explained by the well-documented strong sensitivity of Z-turn tetraloops to their neighboring base pairs ^39^. Together, these data suggest that the non-canonical pairs at pri-miRNA apical junctions contribute to their structural stability in solution.

### U-U and G-A pairs are enriched at apical junctions of human miR-precursors

We next estimated the abundance of non-canonical pairs at human miR-precursor apical junctions. Among 1,881 miR-precursor sequences, 341 contain U residues at both 5’ and 3’ ends that are most likely to pair like in the pri-miR-340, pri-miR-300, and pri-miR-449c structures (Fig. 4e). The U-U pair is the most abundant among all possible combinations at these positions, whereas the expected occurrence by chance is 181 (Fig. 4f). This enrichment is highly significant, as the probability of observing U-U 340 times by chance is 3 × 10^−28^ times lower than that for 181 times. The second most abundant combination is 5’-G and 3’-A, observed 245 times, 1 × 10^−16^ fold less likely to occur by chance than the odd for most probable count of 139. The loop sequence counts of other terminal combinations such as C-A (observed 122 times) are less substantially different from that expected by chance (109 times, *P*_122_/*P*_109_ = 0.42). Therefore, we conclude that human miR-precursors favor U-U and G-A pairs immediately next to the hairpin stem.

Intriguingly, U-U and G-A are known to stabilize hairpin loops when serving as the closing pairs ^40^. Other non-canonical pairs, such as G-G, C-A and A-C, are also known to be stabilizing but to less extents; and they are not significantly enriched in miR-precursor apical junctions. We speculate that U-U and G-A non-canonical pairs are favored by miR-precursor apical junctions for their stabilizing effects and/or specific geometric features.

Non-canonical pairs shorten apical loops by 2 nt and extend hairpin stems by 1 bp. Both properties are key characteristics for miR-precursors ^4, 5, 6, 7^. Therefore, we analyzed the correlation between U-U/G-A pairs and apical loop length by calculating loop length distributions for the corresponding pri-miRNAs in humans. When all pri-miRNAs are included in the analysis, we observed an enrichment of UU and GA pairs among pri-miRNAs with apical loop length between 8 and 16 nt, accompanied by depletion among pri-miRNAs with <7 nt loops (Fig. 4g, Supplementary Fig. 4a,b). Since the authenticity of some pri-miRNAs is debatable ^41^, we also analyzed a set of 319 “empirical” pri-miRNAs that have been previously characterized and enrich well-processed pri-miRNAs ^6^. U-U and G-A pairs are even more pronouncedly enriched among empirical pri-miRNAs with >8-nt loops (Fig. 4h, Supplementary Fig. 4c,d). We suggest that pri-miRNAs with longer apical loops can better tolerate the loop shortening effect of U-U and G-A pairs. Conversely, it is also possible that U-U or G-A pair benefits pri-miRNAs with longer apical loops (supporting evidence presented in the next section). We did not observe statistically significant correlation between U-U/G-A pairs and pri-miRNA stem length, partially due to the fact that pri-miRNA stems are long (optimally 35 bp) with wide dispersion.

### U-U pairs at the apical junction enhance miRNA maturation in human cells

Our structural, biochemical, and sequence analyses suggest that non-canonical pairs at the apical junction may be functionally important. As residues involved in these non-canonical pairs are typically not part of the mature miRNA strands, they are likely to influence miRNA maturation. To test this hypothesis, we employed a cellular miRNA maturation assay based on a bicistronic pri-miRNA expression cassette (Fig. 5a). Using transient transfection in mammalian cells, we expressed transcripts consisting of two fused pri-miRNA sequences. The 3’ pri-miRNA is the subject of interrogation, whereas the 5’ pri-miR-9-1 serves as normalization. Each pri-miRNA sequence is 150-nt in length and contains the hairpin and 33-34 nt flanking sequences on both sides. We measured the abundance of the two mature miRNAs produced from the bicistronic transcripts using quantitative RT-PCR. We found that in HEK293T cells, pri-miR-300 is poorly processed, as indicated by much larger C_t_ values of miR-300 than those for miR-9 (ΔC_t_ around 9). This can be explained by its poorly paired hairpin stem. pri-miR-340 is moderately processed, with miR-340 C_t_ values greater than those of miR-9 by about 1.5. This result makes sense considering the fact that pri-miR-340 has a 40-nt stem and a 7-nt apical loop, just outside the respective 33-39 nt and 8-16 nt ranges deemed optimal for processing (Fig. 5b) ^4, 5, 6^.

**Fig. 5.**
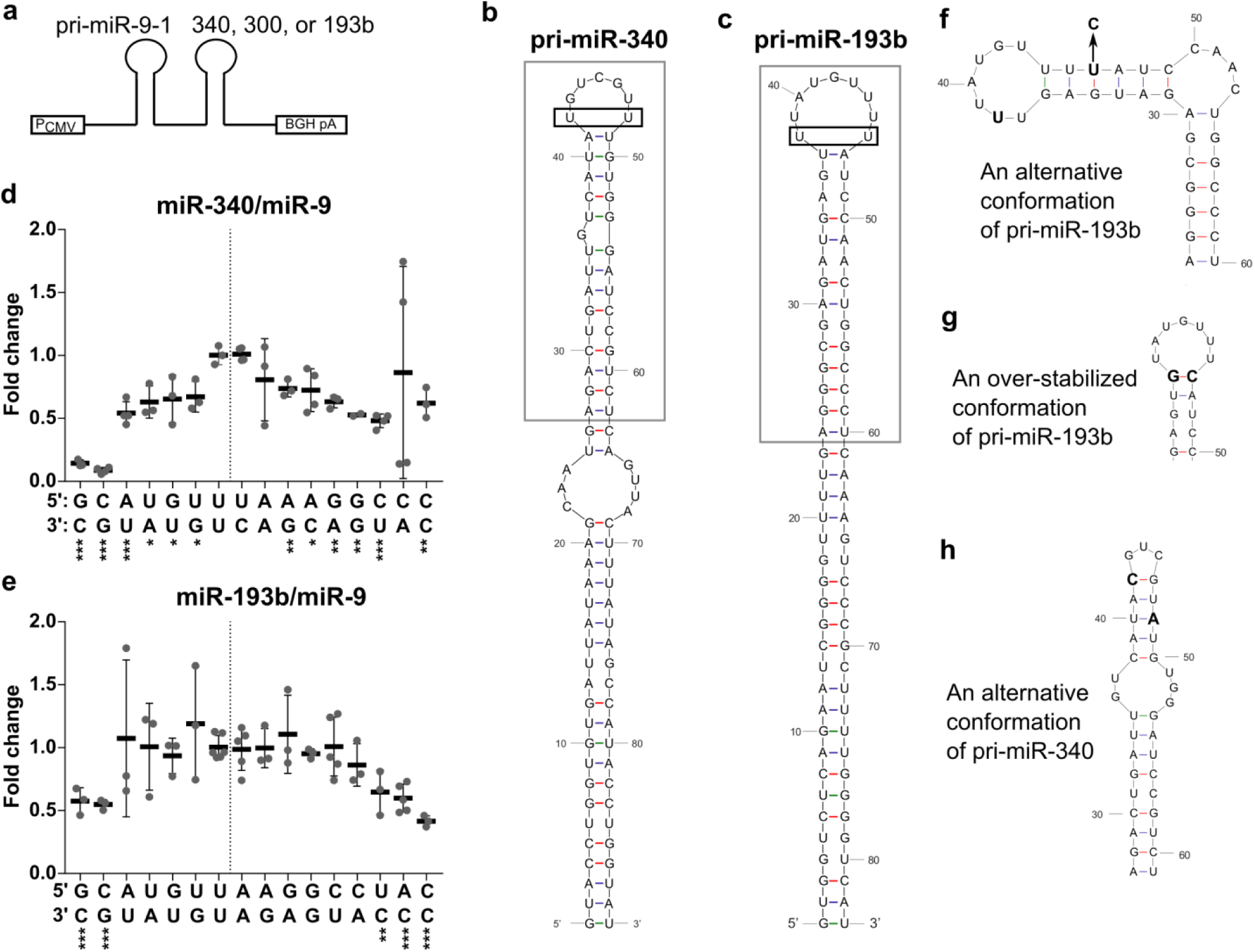
Terminal residues in pri-miRNA apical loops fine-tune miRNA production. **a** Schematic of dual-pri-miRNA constructs for measuring miRNA maturation efficiency in mammalian cells. Each pri-miRNA fragment contains the hairpin and about 30-nt flanking sequence on each side, totaling ∼150 nt. The pri-miR-9-1 fragment is unchanged and is used for normalization. The terminal loop residues of the 3’ pri-miRNA fragment are subjected to mutagenesis. The abundance of both mature miRNAs is measured using quantitative RT-PCR. **b**,**c** Secondary structure of pri-miR-340 and pri-miR-193b hairpin predicted by MFOLD ^34^. The loop-closing UU residues are highlighted in black boxes. Residues flanking the hairpins are not shown. **d** Maturation efficiencies of pri-miR-340 variants (miR-340/miR-9 ratios). **e** Maturation efficiencies of pri-miR-193b variants. In these scatter plots, individual data points are shown as gray dots. The thick bars indicate means and error bars represent standard deviations. Asterisks indicate the P value of the corresponding variant compared to UU, the WT (***, P≤0.001; **, 0.001<P≤0.01; *, 0.01<P≤0.05). The vertical lines divide the variants into two groups, as discussed in the text. **f** A less stable alternative structure of pri-miR-193b is stabilized by a U-to-C mutation. The structure corresponds to the region highlighted by the gray box in **c. g** Changing UU to GC in pri-miR-193b (highlighted in bold) extends the hairpin stem and shortens the apical loop. **h** Changing UU to CA in pri-miR-340 is predicted by MFOLD to alter the secondary structure of the apical region. The structure corresponds to the region highlighted by the gray box in **b**.

We then mutated the U-U pair to all other possible base combinations, performed the cellular miRNA maturation assay. We found that for pri-miR-340, the wild-type sequence, with U-U pair at the apical junction, is the most efficiently processed as compared to other mutants (Fig. 5d). So is U-C, another combination of two pyrimidines that can pair with two hydrogen bonds. In contrast, all canonical RNA base pairs at this position result in much less efficient processing. The strongest G-C and C-G pairs are the least efficiently processed, with miR-340 levels at 14% ± 3% (mean ± SD) and 9% ± 2% of the wild type, respectively. The variants with less stable canonical pairs produce medium levels of miR-340, 54% ± 9% for A-U, 63% ± 13% for U-A, 65% ± 19% for G-U, and 67% ± 12% for U-G. These observations indicate that pairing is not sufficient to enhance miRNA maturation. Rather, the U-U pair may be favored because of its geometry. The miR-340 abundance of most other variants are 48%-74% of U-U with P values lower than 0.05; among them is the G-A pair (63% ± 5%). AA and CA also produce less miR-340, at 81% and 86% of U-U, but large experimental variations prevent the reductions from reaching statistical significance. For the pri-miR-300 variants, low abundance of miR-300 makes the data so noisy that no significant differences can be inferred. These data suggest that a U-U pair at the apical junction serves to expedite miRNA maturation, but is insufficient for rescuing a poorly processed pri-miRNA.

To test if non-canonical pairs are favored for maturation of other pri-miRNAs, we selected pri-miR-193b for similar mutagenesis and cellular miRNA maturation assays. Pri-miR-193b contains a 37-nt stem and a 9-nt apical loop (Fig. 5c). The apical loop is closed by two U residues, which presumably form the same non-canonical pair as in our structures. We found that for wild-type pri-miR-193b, the C_t_ values of miR-193b are smaller than that of miR-9 by about 1.2, indicating that pri-miR-193b is efficiently processed. This observation is consistent with the optimal stem and loop lengths. Among all variants, the average abundance of miR-193b is within a 3-fold range (Fig. 5e). Similar to pri-miR-340, the wild-type pri-miR-193b is among the most efficiently processed. This group of most efficient processors is relatively large, containing 10 mutants with miR-193b abundance within 20% of the wild type. The differences among these most efficient variants are not statistically significant. The 10 mutants include combinations that can form either canonical (A-U, U-A, G-U, and U-G) or non-canonical (A-G, G-A, C-U, and C-A) base pairs, including potential Hoogsteen pairs (A*A and G*G). Also similar to pri-miR-340, the strong G-C and C-G pairs are among the least efficiently processed, with miR-193b levels at 58% ± 11% and 55% ± 4% respectively. The abundance of mature miR-193b for the U-C, A-C, and C-C mutants are significantly lower, at 65% ± 18%, 60% ± 11%, and 42% ± 4% of the wild type respectively. Therefore, unlike pri-miR-340, U-C is not the most efficiently processed for pri-miR-193b (an explanation offered in the next paragraph). We note that the second group of less efficient pri-miR-193b mutants contains 3’ cytosine residues, consistent with C being least abundant as either 5’ or 3’ closing residues among human pri-miRNA apical loops (Fig. 4f). Overall, these results support the conclusion that U-U is a favorable pair for pri-miRNA processing. Our study establishes a function of non-canonical pairs at pri-miRNA apical junctions as fine-tuning miRNA production.

### Correlation between Rhed-binding affinity and apical loop length

We and Kim’s group have previously shown that Rhed binds pri-miRNA apical junction regions ^17, 24, 25^. We wondered whether the Rhed has any preferences for pri-miRNAs with certain sequences, structures and loop lengths. We addressed this question by measuring the affinities of Rhed for fragments of the eight pri-miRNAs containing the apical loop plus approximately 20 bp from the stem (Supplementary Fig. 5). Our electrophoresis mobility shift assays (EMSA) indicated predominantly a single shifted band (Supplementary Fig. 6), which we interpret as a dimeric Rhed protein bound to one RNA molecule. Dissociation constants (*K*_d_) obtained from quantification and fitting range from 1.9 to 9.2 μM (Fig. 6). Such differences could be important for the recognition, especially when pri-miRNAs compete for the processing machinery. Pri-miR-340, which contains the UGU motif at the 5’ side of the loop, binds the Rhed with similar affinity (*K*_d_ = 3.5 µM) to other constructs lacking this sequence. There is no obvious correlation between the presence of non-canonical pairs and Rhed affinity. However, when we plotted the free energy of binding (ΔG_binding_) versus the overall loop length (Fig. 6i), we noticed a trend toward lower ΔG and thus tighter binding for longer loops. This trend became more obvious when we corrected the loop length based on our 3D structures by reducing the length by the number of residues involved in non-canonical pairs (Fig. 6j). We did not count the U-G pair in pri-miR-19b-2 loop due to the loop’s instability (Fig. 4d). The differences in ΔG_binding_ to Rhed among pri-miRNAs tested are within 1 kcal/mol, in the same order of *RT* value (0.62 kcal/mol) at 37°C and thereby sufficient to make substantial differences to the interaction with Microprocessor and processing efficiency, especially when Microprocessor becomes limiting (in many cancer cells for example). Although in this experiment we cannot rule out the possibility that differences in the pri-miRNA stems also contribute to the Rhed affinities, our results suggest that the preferential processing of pri-miRNAs with longer apical loops is mediated the Rhed domain in DGCR8.

**Fig. 6.**
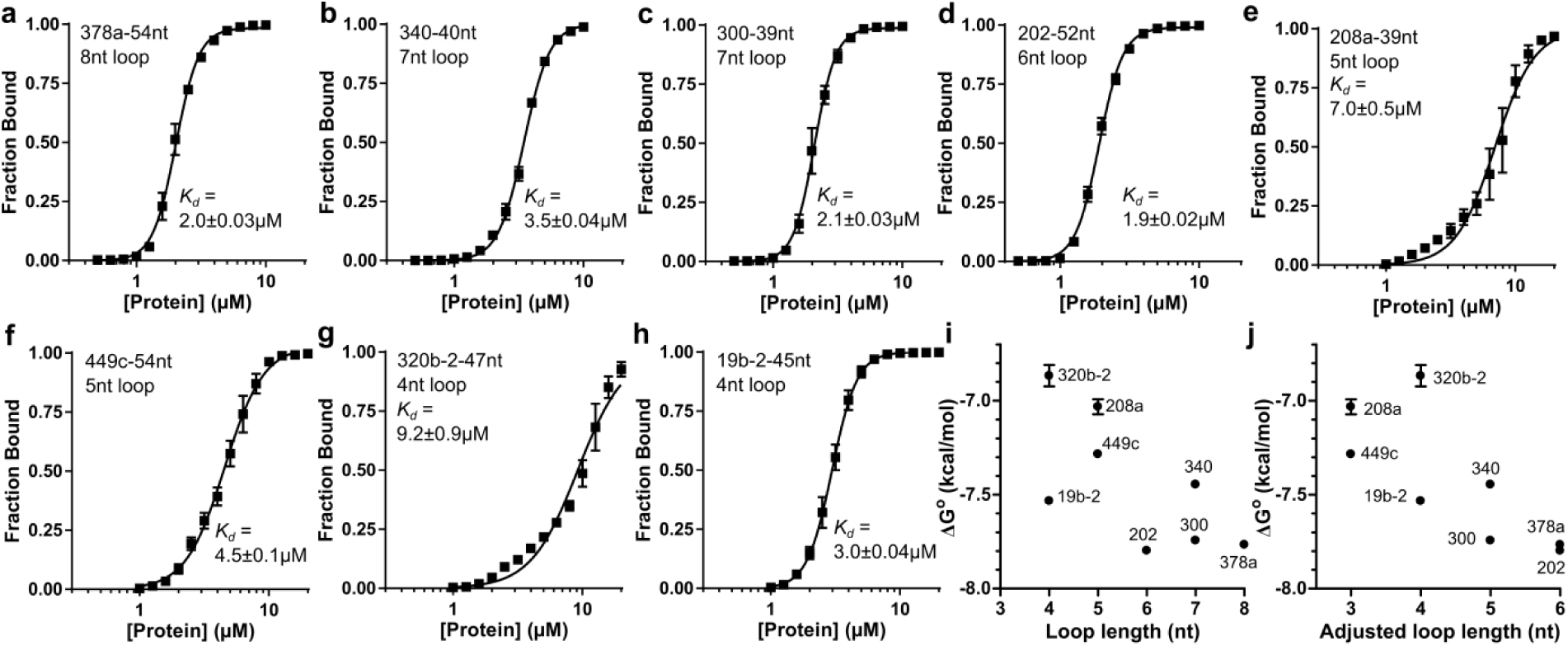
Association of the DGCR8 Rhed domain with pri-miRNA apical junctions. **a**-**h** Quantification of gel shift assays, with representative gel images shown in Supplementary Fig. 6. Data points represent the mean fraction bound ± standard error (SE) from three replicate experiments. Data were fit with the Hill equation and the dissociation constants (*K*_d_) are shown (± SE). **i** Comparison of the free energy of Rhed binding (RTln(*K*_d_)) to the length of the terminal loop, as predicted by mfold. **j** Same as **i** except that the with bases involved in non-canonical pairs excluded. The length of the 19b-2 tetraloop is not adjusted because of its unusually low stability.

## Discussion

We demonstrate scaffold-directed crystallography as a powerful tool for RNA structural biology. In this method, a target RNA is fused to the YdaO scaffold so that it is positioned toward a large solvent cavity in the crystal. As such the target moiety does not disrupt existing lattice contacts, allowing the fusion molecule to be crystalized under conditions very similar to the original. Since rescreening of a broad array of conditions is unnecessary, a minimal amount of purified fusion RNA is required for crystallization. Importantly, because the target does not interact with neighboring molecules in the lattice, its structure represents the conformation in solution. The latter feature is reflected in electron density maps. Namely residues with high structural stability have clear density, whereas the density for more dynamic residues is more defused, and the most flexible residues do not have any density. It is unlikely that the variation in electron density for the target RNAs is an intrinsic property of fusion with the scaffold, as all residues in 378a (the longest loop in our series of structures) and the GAAA tetraloop in WT YdaO are well resolved.

Our method has analogy to the carrier fusion technique often used in protein crystallography, in which a target protein is fused to a carrier protein such as the maltose-binding protein (MBP) via a continuous α-helical linker ^42^. A major problem with the protein scaffolding methods is that the flexibility between the target and the scaffold often results in poorly resolved electron density for the target moieties. Fortunately, in our method the target RNA is fused to the scaffold via a double-stranded helix that is much more rigid than the single α-helical linker in fusion proteins.

The success we report here is limited to short RNA sequences up to 10 nt. This is in part because further extension of the P2 stem may cause the RNA to contact other molecules in the lattice. Longer loops can also interfere with YdaO folding. However, the cavity in the scaffold lattice is large enough to accommodate much more materials, so our method has the potential to reveal larger structures.

Application of scaffold-directed crystallography provides an atomic survey of eight miR-precursor apical junction and loop structures. These structures collectively uncover a structural consensus that involves a non-canonical base pair closing the apical loop and further base stacking at the 5’ end. This consensus is supported by the previously reported NMR structure for pre-miR-20b ^32^. The pre-miR-20b stem terminates in a G-U pair with the neighboring 5’ loop G nucleotide stacks on top (Supplementary Fig. 7). Comparison of the top 20 NMR solutions confirms these as stable features of the molecule. NMR study of pre-miR-21 revealed weak signals corresponding to two tandem U-G/G-U pairs at the apical junction whereas the 14-nt apical loop is otherwise unstructured ^31^.

We asked whether the loop conformations we uncovered were unique to pri-miRNAs or shared with other RNA stem-loops. To address this question, we threaded RNA hairpin sequences from the PDB onto our pri-miRNA structures and then calculated the RMSD between the threaded pose and the original PDB conformation (see Methods). For pri-miR-378a we identified three loops that are slightly shorter (6- or 7-nt) and differ in sequence but retain a highly similar fold (Supplementary Fig. 8). Comparing these structures reveals a generalized loop motif, which we call 3’-purine-rich stack (Supplementary Fig. 8b). In a 3’-purine-rich stack, 4-5 mostly purine bases on the 3’ side of the loop stack with each other, on top of the helical stem. One or two pyrimidines may be found in positions furthest from the stem. On the 5’ side of the loop, two or three pyrimidine residues, most often uridines, serve as linkers between the 3’ stacked residues and the stem. These linker pyrimidines form hydrogen bonds with stacked purines, sometimes non-canonical base pairs, which further stabilize the loop. Many pri-miRNA and other hairpin loops contain sequences consistent with a 3’ purine stack. Furthermore, the AAGU loop in pri-miR-320a is found in Rnt1 substrates in Saccharomyces cerevisiae (an organism that does not encode miRNAs) and the Z-turn formed by the pri-miR-19b-2 tetraloop is commonly found in ribosomal RNAs. Overall, these observations suggest that the pri-miRNA loop structures are not necessarily unique to pri-miRNAs.

We found that a U-U pair at the apical junction is among the most efficiently processed variants of pri-miR-340 and pri-miR-193b (Fig. 5d,e). It is possible that the U-U pair presents favorable geometry to the processing machinery. We would like to offer a thermodynamic explanation for why non-canonical pairs are favored at pri-miRNA apical junctions. Non-canonical pairs balance needs to stabilize the processing-competent conformation, to prevent alternative conformations, and to avoid over-stabilizing and over-extending hairpin stem. This idea is best illustrated using pri-miR-193b as an example. In addition to the processing-competent conformation shown in Fig. 5e, MFOLD predicts an alternative conformation that is not expected to be compatible for processing (Fig. 5f). The alternative conformation is less stable than the WT, with the folding free energy ΔG higher by 0.6 kcal/mol, and thereby is unlikely to strongly impact processing. In the UC, AC, and CC mutants, however, changing the 3’-U to a C introduce a G-C pair that selectively stabilizes the alternative conformation (Fig. 5f), making it the only dominant species. On the other hand, replacing the U-U pair with either a G-C or C-G pair does stabilize the hairpin conformation but also extends the stem by one base pair and reduces the loop by two nucleotides (Fig. 5g). The latter changes move the hairpin structure away from optimal and not surprisingly reduce processing efficiency. Consistent with our theory, a pri-miR-340 variant (CA) is also predicted to adopt an alternative conformation that is incompatible with processing (Fig. 5h). Therefore, we suggest that a function of non-canonical pairs at the apical junction is to stabilize processing-compatible conformation and suppress processing-incompatible alternative conformations. We call this explanation the conformation dynamics theory. Under this theory, since the stability of other canonical pairs, including A-U, U-A, G-U, and U-G, are in between non-canonical pairs and G-C/C-G pairs, their effects on processing may also be somewhere in between. As RNA residues have high tendency of forming base pairs, we believe that alternative conformations are abundant in pri-miRNA apical loops and thereby this function of non-canonical pairs is likely to be general.

Both favorable geometry and conformation dynamics theories are supported by phylogenetic evidence. The U-U pair in pri-miR-340 is nearly completely conserved, whereas nucleotide variation occurs in all other positions. The only variation of the U-U pair is a substitution by a U-G pair in black fruit bat (*Pteropus alecto*). The U-U pair in pri-miR-193b is also conserved among 32 homologs collected in miRBase ^29^, with higher degree of variation than pri-miR-340. In armadillo (*Dasypus novemcinctus*), domestic dog (*Canis familiaris*), spider (*Parasteatoda tepidariorum*), the 9-nt apical loop in humans is changed to 10, 7, 4 nt respectively but the UU pair is maintained, supporting the favorable geometry theory. In zebrafish (*Danio rerio*) and lancelets (*Branchiostoma belcheri*) UU is replaced by a canonical AU pair and in acorn worm (*Saccoglossus kowalevskii*) UU is replaced by AC. These substitutions may be viewed as acceptable under the conformation dynamics theory. Lastly, we note that the two hypotheses are not mutually exclusive, as miRNA maturation includes multiple steps and a myriad of factors serve core and regulatory functions.

Microprocessor recognizes a pri-miRNA hairpin by clamping its stem at both ends ^15, 17^. The optimal pri-miRNA hairpin stem length is estimated to be 35 ± 1 bp, counting in internal non-canonical pairs ^5^. Our study suggests that terminal non-canonical pairs at apical junctions have to be considered. The pri-miR-340 and pri-miR-193b subjected to systematic mutagenesis analyses represent long and near-optimal stem lengths. We imagine that in cases where pri-miRNA helical stems are shorter than optimal, terminal non-canonical pairs at apical and basal junctions could help the hairpin fits in the Microprocessor complex and facilitate their maturation.

## Methods

### Key resources table

| **Reagent type** | **Designation** | **Source or reference** | **Identifiers** | **Additional information** |
| --- | --- | --- | --- | --- |
| recombinant DNA reagent | YdaO-WT-pUC19 (plasmid) | this paper |  | transcription template |
| recombinant DNA reagent | YdaO-pri-miR-9-1-0bp-pUC19 (plasmid) | this paper |  | transcription template |
| recombinant DNA reagent | YdaO-pri-miR-9-1-1bp-pUC19 (plasmid) | this paper |  | transcription template |
| recombinant DNA reagent | YdaO-pri-miR-9-1-2bp-pUC19 (plasmid) | this paper |  | transcription template |
| recombinant DNA reagent | YdaO-pri-miR-9-1-3bp-pUC19 (plasmid) | this paper |  | transcription template |
| chemical compound, drug | c-di-AMP | Sigma | Cat#SML1231 |  |
| Peptide, recombinant protein | BamHI-HF restriction endonuclease | New England Biolabs | Cat#R3136S |  |
| Peptide, recombinant protein | Inorganic pyrophosphatase | Sigma-Aldrich | Cat#I1643 |  |
| Peptide, recombinant protein | T7 RNA polymerase | This study | Prepared in lab |  |
| Peptide, recombinant protein | Inorganic pyrophosphatase | Sigma-Aldrich | Cat#I1643 |  |
| Peptide, recombinant protein | Q5 high-fidelity DNA polymerase | New England Biolabs | Cat#M0530L |  |
| Peptide, recombinant protein | Phusion high-fidelity DNA polymerase | Thermo Fisher Scientific | Cat# F530L |  |
| Software, algorithm | Prism (version 7) | GraphPad Software | https://www.graphpad.com |  |
| Software, algorithm | PyMOL | Schrödinger, LLC | https://pymol.org |  |
| Software, algorithm | XDS | (Kabsch, 2010) | http://xds.mpimf-heidelberg.mpg.de/ |  |
| Software, algorithm | PHENIX | (Adams et al., 2010) | https://www.phenix-online.org/ |  |
| Software, algorithm | Coot | (Emsley et al., 2010) | https://www2.mrc-lmb.cam.ac.uk/personal/pemsley/coot/ |  |
| Software, algorithm | Rosetta | (Cheng et al., 2015) | https://www.rosettacommons.org/ |  |
| Software, algorithm | GROMACS | (Abraham et al., 2015) | http://www.gromacs.org/ |  |
| Software, algorithm | Quantity One | Bio-Rad |  |  |
| Software, algorithm | Bowtie2 | (Langmead and Salzberg, 2012) |  |  |

### Pri-miRNA secondary structure prediction and apical loop sequence analysis

We downloaded from miRBase (release 21) all annotated human miRNA hairpin sequences and their genomic coordinates. The hairpins typically include the pre-miRNA moiety along with a variable number of additional base pairs from the basal stem. For each hairpin, we used the genomic sequence to extend the RNA an equal number of nucleotides at the 5’ and 3’ ends until the total length equaled 150 nt. Thus, this 150-nt window also takes into account the single-stranded RNA on either side of the hairpin. We then generated predicted secondary structures for all pri-miRNA hairpins using mfold ^34^, and generally retained the structures with the lowest predicted free energy of folding.

We manually reviewed all the predictions to ensure they reflected the expected hairpin structure with mature miRNA sequences derived from either or both strands of the stem; in cases where mfold predicted alternative conformations, we selected the structure with the lowest free energy that contained a stem length of approximately three helical turns. We also compared the secondary structures with those from the miRBase. We eliminated 1-2 base pairs in the hairpin that are isolated from the stem and thereby deemed to be unstable. From the compiled list of pri-miRNA apical loops, we extracted the length (*L*) distribution using Excel (Microsoft, Seattle, WA) and plotted the distribution using Prism (GraphPad, La Jolla, CA) (Fig. 1a).

The occurrences of single nucleotides and combinations at terminal positions of the loops were calculated using Excel and plotted using Prism (Fig. 4e,f,g,h). The expected count of each combination was calculated as 1,881 (total number of sequences) multiplied by the probability of observing a nucleotide pair (*P*_*UU*_ for U residues at both first and last positions for example) at random (the null hypothesis) as *P*_*UU*_ = *F*_*U*,1_ × *F*_*U,L*_, where *F*_*U*,1_ and *F*_*U,L*_ are the frequencies of U at loop positions 1 and *L* respectively. The probability of observing UU combination n times out of 1,881 sequences randomly is 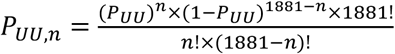. We determined whether a combination is enriched among pri-miRNA apical loops by calculating the ratio of the probabilities of observed UU count over the expected count.

### PDB mining and identification of YdaO crystallization scaffold

We first filtered the PDB to obtain X-ray structures containing RNA but no protein or DNA molecules. To identify voids in the crystal lattices, we wrote a PyMOL script that implemented a grid search algorithm in the following steps. (a) Generate a 3 × 3 × 3 block of unit cells (i.e. 27 copies of the unit cell). The unit cell at the center of this block sees all possible lattice voids, either internally or between unit cells. (b) Using three unit vectors along each of the unit cell axis (i.e. *a, b*, and *c* vectors of length 1 Å), iteratively generate grid points within the central unit cell in the form of 5*i\**a* + 5*j\**b* + 5*k\**c*, where i,j,k are integers. This gives grid points with 5 Å spacing along each axis. (c) For each grid point, calculate the distances to all C1’ atoms in the super cell and identify the shortest as R_local_. For each structure, identify the grid point with the largest R_local_ as R_max_.

To find suitable scaffolds, we then manually reviewed the structures with large R_max_ values, focusing on those with a single RNA chain in the asymmetric unit but also including select candidates with multiple chains (Fig. 1b). We traced the chain looking for any stem-loop that projected into the cavity in the lattice. Amongst several hundred candidates reviewed, only the P2 stem-loop from the YdaO riboswitch (PDB ID: 4QK8) met these conditions ^35^.

### Preparation of YdaO WT and pri-miR-9-1 fusion RNAs and native gel electrophoresis

We initially designed the WT YdaO construct to contain a T7 promoter sequence at the 5’ end and HDV ribozyme on the 3’ side, along with flanking EcoRI and BamHI restriction sites. This fragment was synthesized as a gene block (Integrated DNA Technologies, Coralville, IA), double digested and cloned into the pUC19 plasmid. The clone was verified by Sanger sequencing. To replace the P2 loop nucleotides with the pri-miRNA stem-loop, we used a two-round PCR protocol. All reactions were performed with Q5 high-fidelity DNA polymerase (New England Biolabs, Ipswich, MA) following the manufacture’s recommended reaction setup and cycling conditions. All reactions contained the same reverse primer, which annealed to the 3’ end of HDV and contained the BamHI site (5’-CGT<u>GGATCC</u>GGTCCCATTC-3’). For the first PCR, the forward primer contained the pri-miRNA sequence (underlined) plus around 20 nt upstream and downstream sequences. The forward primers for pri-miR-9-1 fusions were 5’-*CTATA***GGTTGCCGAATCC**<u>GTGGTGTGGAGTCT</u>**GGTACGGAGGAACCGCTTTTTG**-3’ (pri-miR-9-1+0bp), 5’-*CTATA***GGTTGCCGAATCC**<u>AGTGGTGTGGAGTCTT</u>**GGTACGGAGGAACCGCTTTTTG**-3’ (pri-miR-9-1+1bp), 5’-*CTATA***GGTTGCCGAATCC**<u>GAGTGGTGTGGAGTCTTC</u>**GGTACGGAGGAACCGCTTTTTG**-3’ (pri-miR-9-1+2bp), and 5’-*CTATA***GGTTGCCGAATCC**<u>AGAGTGGTGTGGAGTCTTCT</u>**GGTACGGAGGAACCGCTTTTTG**-3’ (pri-miR-9-1+3bp). The scaffold residues are shown in bold. This PCR product was gel-purified and 1 μL was used as template for the second-round PCR. All second-round reactions contained the same reverse primer and a forward primer (5’-GCA<u>GAATTC</u>*TAATACGACTCACTATA***GGTTGCCGAATCC**-3’), which added the full T7-promoter (italic) and EcoRI site (underlined). The second-round PCR product was gel-purified, digested with EcoRI and BamHI, and ligated into pUC19. Clones containing the desired insert were sequence-verified.

For WT YdaO and pri-miR-9-1 fusion constructs we prepared maxiprep plasmids and linearized them by overnight digestion with BamHI. Transcription reactions contained ∼400 μg (45 nM) linearized template, 40 mM Tris pH7.5, 25 mM MgCl_2_, 4 mM DTT, 2 mM spermidine, 40 μg inorganic pyrophosphatase (Sigma, St. Louis, MO), 0.7 mg T7 RNA polymerase, and 3 mM each NTP in a total volume of 5 mL. After 4.5 hr of incubation at 37°C, the final MgCl_2_ concentration was adjusted to 40 mM, and the reactions were incubated for additional 45 min. Despite the elevated Mg^2+^ concentration, we observed only partial cleavage by the HDV ribozyme. Reactions were ethanol precipitated and purified over denaturing 10% polyacrylamide slab gels. The desired product was visualized by UV shadowing and excised from the gel. Gel pieces were crushed and extracted overnight in 30 mL TEN buffer (150 mM NaCl, 20 mM Tris pH 7.5, 1 mM EDTA) at 4°C. We then spun down the gel pieces and concentrated the RNA in an Amicon Ultra-15 centrifugal filter unit with 10-kDa molecular weight cutoff (MWCO). RNA was buffer-exchanged three times into 10 mM NaHEPES pH 7.5 and concentrated to ∼50 μL final volume.

For analysis on a native gel, 5 μM RNA stock solutions were prepared by dilution of the purified RNA into 5 mM Tris pH 7.0. Next, 2.5 μL RNA was mixed with an equal volume of 2X annealing buffer containing 35 mM Tris pH 7.0, 100 mM KCl, 10 mM MgCl_2_, and 20 μM c-di-AMP (Sigma). The mixtures were heated at 90°C for 1 min followed by snap cooling on ice and then a 15-min incubation at 37°C. The annealed RNA was mixed with a 2X loading dye containing 40 mM Tris pH 7.0, 50 mM KCl, 5 mM MgCl_2_, 20% (v/v) glycerol, and xylene cyanol, and analyzed at 4°C on a 10% polyacrylamide gel with Tris-borate (TB) running buffer. The gel was stained in Sybr Green II and scanned on a Typhoon 9410 Variable Mode Imager (GE Healthcare, Piscataway, NJ).

### Preparation of pri-miRNA-YdaO fusions for crystallization

Given the poor HDV self-cleavage efficiency we observed for the pri-miR-9-1 fusions, we elected to change strategy. Instead of employing a ribozyme to create homogeneous 3’ ends, we used PCR to generate transcription templates in which the two 5’ residues on the anti-sense DNA strand were 2’-O-methylated. The modifications have been shown to reduce un-templated nucleotide addition by T7 RNA polymerase ^43^. We utilized a three-round PCR approach to create the transcription templates. All reactions below contained the same reverse primer, 5’-mCmUCCTTCCTTTATTGCCTCC-3’, where ‘m’ indicates 2’-O-methylation. For the first round of PCR, we set up a 50 μL reaction with Q5 polymerase to amplify the 3’ fragment of YdaO with the forward primer 5’-GGTACGGAGGAACCGCTTTTTG-3’ and performed 30 cycles of amplification. The product was gel purified and 1 μL was used as template for the next round. In the second-round PCR, we used a unique forward primer for each construct containing the pri-miRNA loop and stem sequence which annealed to the 3’ YdaO fragment from the first stage. The primer sequences were 5’-*CTATA***GGTTGCCGAATCC**<u>ATATGT</u>**GGTACGGAGGAACCGCTTTTTG**-3’ (19b-2+1bp), 5’-*CTATA***GGTTGCCGAATCC**<u>GATCTGGC</u>**GGTACGGAGGAACCGCTTTTTG**-3’ (202+1bp), 5’-*CTATA***GGTTGCCGAATCC**<u>GATGCTC</u>**GGTACGGAGGAACCGCTTTTTG**-3’ (208a+1bp), 5’-*CTATA***GGTTGCCGAATCC**<u>CTTTACTTG</u>**GGTACGGAGGAACCGCTTTTTG**-3’ (300+1bp), 5’-*CTATA***GGTTGCCGAATCC**<u>AAAGTT</u>**GGTACGGAGGAACCGCTTTTTG**-3’ (320b-2+1bp), 5’-*CTATA***GGTTGCCGAATCC**<u>ATGTCGTTT</u>**GGTACGGAGGAACCGCTTTTTG**-3’ (340+1bp), 5’-*CTATA***GGTTGCCGAATCC**<u>ACCTAGAAAT</u>**GGTACGGAGGAACCGCTTTTTG**-3’ (378a+1bp), and 5’-*CTATA***GGTTGCCGAATCC**<u>ATGATTT</u>**GGTACGGAGGAACCGCTTTTTG**-3’ (449c+1bp). This reaction was also 50 μL and used Q5 polymerase for 30 cycles. The product from the second-round PCR was analyzed by agarose gel electrophoresis to confirm amplification, and 40 μL of the reaction was used as template for the third-round PCR without further purification. The 2-mL PCR reactions used the Phusion high-fidelity DNA polymerase (Thermo Fisher Scientific, Carlsbad, CA) and the forward primer 5’-GCA<u>GAATTC</u>*TAATACGACTCACTATA***GGTTGCCGAATCC**-3’, and was run for 35 cycles.

The third-stage PCR product was purified over a HiTrap Q HP column (GE Healthcare). Buffer A contained 10 mM NaCl and 10 mM NaHEPES pH 7.5; Buffer B was identical but with 2 M NaCl. All NaHEPES buffers used in this study were prepared by titrating HEPES free acid solution with NaOH. The column was equilibrated with 20% Buffer B and the desired DNA product was eluted with a linear gradient to 50% B over 10 min at 2 ml/min. We analyzed the peak fractions on an agarose gel to confirm they contained a single band of the correct size. The peak fractions were then pooled and concentrated in an Amicon filter unit (10 kDa MWCO), and then washed with water to remove excess salt. The concentration of the DNA template (∼200 μL final volume) was determined by UV absorbance.

Transcription reactions were set up as described above for pri-miR-9-1 fusions, but in a 10-mL volume and containing 28 nM DNA template. Reactions were run for 4 hr at 37°C followed by phenol-chloroform extraction. The transcription was concentrated in an Amicon filter unit (10 kDa MWCO) and washed with 0.1 M trimethylamine-acetic acid (TEAA) pH 7.0. The RNA (∼2 mL) was injected onto a Waters XTerra MS C18 reverse phase HPLC column (3.5 μm particle size, 4.6 × 150 mm in dimension) thermostated at 54°C. TEAA and 100% acetonitrile were used as mobile phases. The column was washed with 6% acetonitrile and the RNA eluted with a gradient to 17% acetonitrile over 80 min at 0.4 ml/min. Peak fractions were analyzed on denaturing 10% polyacrylamide gels. Pure fractions were pooled and buffer-exchanged into 10 mM NaHEPES pH 7.0 using an Amicon filter unit. The RNA was concentrated to <50 μL final volume and the concentration determined by UV absorbance.

### Crystallization, data collection, and structure determination

All RNA-c-diAMP complexes were prepared as described ^35^. Briefly, a solution containing 0.5 mM RNA, 1 mM c-di-AMP, 100 mM KCl, 10 mM MgCl_2_, and 20 mM NaHEPES pH 7.0 was heated to 90°C for 1 min, snap cooled on ice, and equilibrated for 15 min at 37°C immediately prior to crystallization. Screening was performed in 24-well plates containing 0.5 mL well solution; the hanging drops consisted of 1 μL RNA plus 1 μL well solution. Plates were incubated at room temperature, and crystals generally grew to full size (100 μm to over 200 μm) within one week. For 19b-2+1bp, the well solution contained 1.7 M (NH_4_)_2_SO_4_, 0.2 M Li_2_SO_4_, and 0.1 M NaHEPES pH 7.1. For 202+1bp, 208a+1bp, and 320b-2+1bp, the well contained 1.9 M (NH_4_)_2_SO_4_, 0.2 M Li_2_SO_4_, and 0.1 M NaHEPES pH 7.4. The well solution for 378a+0bp contained 1.7 M (NH_4_)_2_SO_4_, 0.2 M Li_2_SO_4_, and 0.1 M NaHEPES pH 7.4. For the remaining constructs crystallization was performed in 96-well plates with hanging drops consisting of 0.4 μL RNA plus 0.4 μL well solution. For 300+0bp, the well solution contained 1.90 M (NH_4_)_2_SO_4_, 0.158 M Li_2_SO_4_, and 0.1 M NaHEPES pH 7.4. Construct 340+1bp crystallized from a well solution containing 1.89 M (NH_4_)_2_SO_4_, 0.214 M Li_2_SO_4_, and 0.1 M NaHEPES pH 7.4. Construct 378a+1bp crystallized from 1.63 M (NH_4_)_2_SO_4_, 0.272 M Li_2_SO_4_, and 0.1 M NaHEPES pH 7.4. For construct 449c+1bp, the well contained 1.89 M (NH_4_)_2_SO_4_, 0.128 M Li_2_SO_4_, and 0.1 M NaHEPES pH 7.4.

All crystals were briefly soaked in a cryoprotectant solution containing 20% (w/v) PEG 3350, 20% (v/v) glycerol, 0.2 M (NH_4_)_2_SO_4_, 0.2 M Li_2_SO_4_, and 0.1 M NaHEPES pH 7.3, and then flash-frozen in liquid nitrogen. Data were collected at 100 K at the Advanced Photon Source Beamline 24-ID-C or the Advanced Light Source Beamline 8.3.1. For all constructs we collected a native dataset at a wavelength of ∼1 Å. For 378a+0bp, 202+1bp, 449c+1bp, and 320b-2+1bp, we measured phosphorous anomalous scattering by collecting additional high-redundancy datasets at 1.9 Å. Data were indexed, integrated, and scaled using XDS ^44^.

Where anomalous data were available, we generated partially-experimental phases using a combined molecular-replacement/single anomalous dispersion approach (MR-SAD). The molecular replacement model consisted of the YdaO c-di-AMP riboswitch structure (PDB ID: 4QK8) with the GAAA tetraloop on the P2 stem removed from the model. Phases were obtained using the default settings in the Phaser-MR protocol in Phenix ^45^.

For all constructs we obtained an initial solution by performing a rigid-body fit of the MR model (above) to data using Phenix (including experimental phase restraints where available). This produced an excellent initial model with R_work_ < 30%. We then inspected the electron density map in region of the P2 stem. For all RNAs, additional density for the missing base-pair and loop could clearly be seen in the 2F_o_-F_c_ and difference maps. We then modeled in the missing residues in Coot ^46^. In cases where the density was unclear, we stopped modeling with an incomplete loop and performed an additional round of coordinate, ADP, and TLS parameter refinement with Phenix. This typically revealed additional density for the missing residues. As mentioned in the main text, a few residues did not end up having sufficient density for determining their conformations. We completed the loop models by including the most likely conformation for these residues. We performed subsequent rounds of refinement and manual adjustment as above until reasonable R factors and model geometry were obtained.

Sigma A-weighted simulated annealing omit maps were calculated using Phenix (Supplementary Figs. 1 and 2). All pri-miRNA residues were omitted. Simulated annealing was performed in Cartesian space with an annealing temperature of 5000°C. We excluded bulk solvent from omit regions. This type of omit map is known as Polder map and prevents the solvent mask from obscuring relatively weak densities ^47^.

### Optical melting

RNA for optical melting experiments were transcribed *in vitro* from synthetic DNA templates (Integrated DNA Technologies). The oligonucleotide template sequences used were 5’-GGAAC**ACATAT**GTTCC*TATAGTGAGTCGTATTA*-3’ (19b-2), 5’-GGAAC**GCCAGATC**GTTCC*TATAGTGAGTCGTATTA*-3’ (202), 5’-GGAAC**GAGCATC**GTTCC*TATAGTGAGTCGTATTA*-3’ (208a), 5’-GGAAC**CAAGTAAAG**GTTCC*TATAGTGAGTCGTATTA*-3’ (300), 5’-GGAAC**AACTTT**GTTCC*TATAGTGAGTCGTATTA*-3’ (320b-2), 5’-GGAAC**AAACGACAT**GTTCC*TATAGTGAGTCGTATTA*-3’ (340), 5’-GGAAC**ATTTCTAGGT**GTTCC*TATAGTGAGTCGTATTA*-3’ (378a), and 5’-GGAAC**AAATCAT**GTTCC*TATAGTGAGTCGTATTA*-3’ (449c), with the T7 promoter shown in italics and the pri-miRNA junction/loop segment in bold. Templates were annealed with a second strand complementary to T7 promoter and added to large-scale (10 mL) transcription reactions as described above. Reactions were ethanol precipitated, purified over 20% polyacrylamide denaturing gels. The desired band recovered by UV shadowing. Following gel extraction, samples were buffer exchanged into water and concentrated in an Amicon centrifugal filter device.

For each RNA, a set of 6 dilutions were prepared in 50 mM NaCl and 10 mM sodium cacodylate pH 7.0, such that the initial absorbance ranged from ∼1.0 to 0.1 AU. The samples were annealed by heating to 95°C for 1 min and snap cooling on ice, followed by equilibration to 12°C. Melting measurements were performed with a Cary Bio300 UV-visible spectrophotometer equipped with a Peltier-type temperature controlled sample changer. The absorbance at 260 nm was recorded while the RNA was heated from 12°C to 92°C at a rate of 0.8°C/min. Melting curves were analyzed using Prism (GraphPad, version 7) and fit with the equation 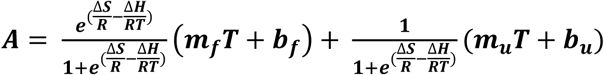, where the absorbance (*A*) is approximated as a function of temperature (T). The changes in entropy (Δ*S*) and enthalpy (Δ*H*) were fit as well as the slope (m) and y-intercept (b) for both the double-stranded (***m***_***f***_ and ***b***_***f***_) and single-stranded (***m***_***u***_ and ***b***_***u***_) linear regions. The melting temperatures and thermodynamic parameters at 37°C were then derived from these parameters (Table 2).

### Cellular miRNA maturation assays

These assays were performed similarly to the report by Nam and colleagues ^24^. Two 150-nt pri-miRNA fragments were juxtaposed together via PCR and were inserted into the pcDNA3.1/Hygro(+) vector between NheI and XbaI sites. The 5’ fragment is pri-miR-9-1, which serves as normalization (Fig. 5a). The 3’ fragment is either pri-miR-340, 300, or 193b, which was subjected to mutagenesis. After realizing that NheI and XbaI have compatible cohesive ends, we switched to use ApaI as the downstream cloning site, but kept the XbaI sequence in the dual-pri-miRNA expression constructs so that their non-pri-miRNA sequences were the same. Mutagenesis was performed using 4-primer PCR. Most mutant pri-miR-340, 300, 193b constructs were generated using primers containing mixed bases at the desired positions. The mutant clones were identified using Sanger sequencing. This strategy of random mutagenesis followed by sequencing usually misses a few clones in each series, which were eventually produced using mutant-specific primers.

The dual pri-miRNA expression plasmids were transfected to human embryonic kidney (HEK) 293 cells in 6-well plates using the Lipofectamine 3000 reagents (Thermo Fisher Scientific) following standard procedures. Forty hours posttransfection, total RNAs were extracted from cells using the TRIZOLE reagent (Thermo Fisher Scientific). The abundance of miRNAs were measured using the Taqman MicroRNA Assays (Thermo Fisher Scientific).

### Electrophoresis mobility shift assay

Human heme-bound Rhed protein was over-expressed in *E. coli* and purified using ion exchange and size exclusion chromatography, as previously described ^17^. Radiolabeled pri-miRNA stem-loops (Supplementary Fig. 5) were prepared by *in vitro* transcription. DNA templates consisted of anti-sense oligonucleotides covering the desired sequence plus the T7 promoter, annealed to a sense oligo with the T7 promoter sequence ^48^. Each 20-μL transcription reaction contained 50 fmol template, 40 mM Tris pH 7.5, 25 mM MgCl_2_, 4 mM DTT, 2 mM spermidine, 2 μg T7 RNA polymerase, 0.5 mM ATP, 3 mM each of UTP, CTP, and GTP, and 3 nmol α-^32^P-ATP (10 μCi). Transcriptions were run at 37°C for 2 hr and the RNA purified over a denaturing 15% polyacrylamide gel. The RNA were extracted overnight at 4°C in TEN buffer, isopropanol-precipitated, and resuspended in 40 μL water.

We adopted a recently reported EMSA procedure to examine Rhed-pri-miRNA interactions ^24^. The RNAs were diluted in 100 mM NaCl, 20 mM Tris pH 8.0 and heated at 90°C for 1 min followed by snap cooling on ice. The annealed RNA was added to binding reactions containing 10% (v/v) glycerol, 0.1 mg/ml yeast tRNA, 0.1 mg/ml BSA, 5 μg/ml heparin, 0.01% (v/v) octylphenoxypolyethoxyethanol (IGEPAL CA-630), 0.25 unit RNase-OUT ribonuclease inhibitor, xylene cyanol, 20 mM Tris pH 8.0, and 0-20 μM Rhed protein. The final salt concentration of the solution was 150 mM NaCl. Binding reactions were incubated at room temperature for 30 min prior to loading on a 10% polyacrylamide gel. Both the gel and the running buffer contained 80 mM NaCl, 89.2 mM Tris base, and 89.0 mM boric acid (pH 8.2 final). Gels were run at 110 V for 45 min at 4°C, and then dried and exposed to a storage phosphor screen. Screens were subsequently scanned on a Typhoon scanner (GE Healthcare). The free and bound RNA bands were quantified using Quantity One software (Bio-Rad, Hercules, CA) and fit with the Hill equation in Prism.

### Comparison to known RNA loop structures in the PDB

To identify RNA loops in the PDB with structural similarity to our pri-miRNA loop models, we first extracted the coordinates for the pri-miRNA apical junctions and loops. The search pool was the same set of RNA structures used to identify crystallization scaffolds above. For each structure from the PDB set, we used DSSR to identify all hairpin loops. We extracted the RNA sequence from each hairpin loop, and eliminated loops shorter than the pri-miRNA sequence. For loops longer than the pri-miRNA, we used a sliding window to obtain all fragments of the loop with the same length. Each loop sequence was then threaded onto the pri-miRNA model using the “rna_thread” routine in Rosetta ^49^. Using a PyMOL script, we aligned the resulting threaded model to the original hairpin loop and calculated the RMSD between the two models. We aggregated and sorted the RMSD data from all PDB structures and manually inspected loops with small RMSD to find hits with structural similarity.

### Statistics

Standard statistical methods were used in crystallographic computing. Probability calculation for apical loop sequence analyses is detailed in its section earlier in Methods. The cellular miRNA maturation assay was independently repeated 3-6 times as indicated in Fig. 5d,e. The P values for the results were calculated in Excel using one-tailed Student’s *t* test assuming 2-sample equal variance.

### Data availability

The coordinates and structure factors have been deposited in PDB under the accession codes indicated in Table 1. All other data are available from the corresponding author upon reasonable request.

## Acknowledgements

We thank Duilio Cascio and Michael Sawaya from the UCLA-DOE Crystallization and X-ray Diffraction facilities for guidance in data collection and access to computing resources. Malcolm Capel and Kay Perry at the Advanced Photon Source sector 24 (NE-CAT) also assisted with data collection; NE-CAT is supported by funding from the NIH (P41 GM103403 and S10 RR029205), and DOE (DE-AC02-06CH11357). We also thank George Meigs and Jane Tanamachi at ALS beamline 8.3.1 for assistance; BL 8.3.1 is supported by the UC Multicampus Research Programs and Initiatives grant (MR-15-328599), and the NIH (R01 GM124149 and P30 GM124169). This project was in part supported by the National Science Foundation (Grant # 1616265 to F.G). G.S. was supported by an NIH/CMB Ruth L. Kirschstein National Research Service Award (GM007185) and the Philip J. Whitcome Fellowship.

## Author contributions

G.M.S. and F.G. conceived the idea of RNA scaffold-directed crystallography and wrote the manuscript. G.M.S. and Z.P. performed the experiments. G.M.S determined the structures with help from Z.P. and F.G. at refinement. All authors jointly did the pri-miRNA loop analyses.

