## Supplemental Figures for "Structures of microRNA-precursor apical junctions and loops reveal non-canonical base pairs important for processing"

*Figure Supplements for*

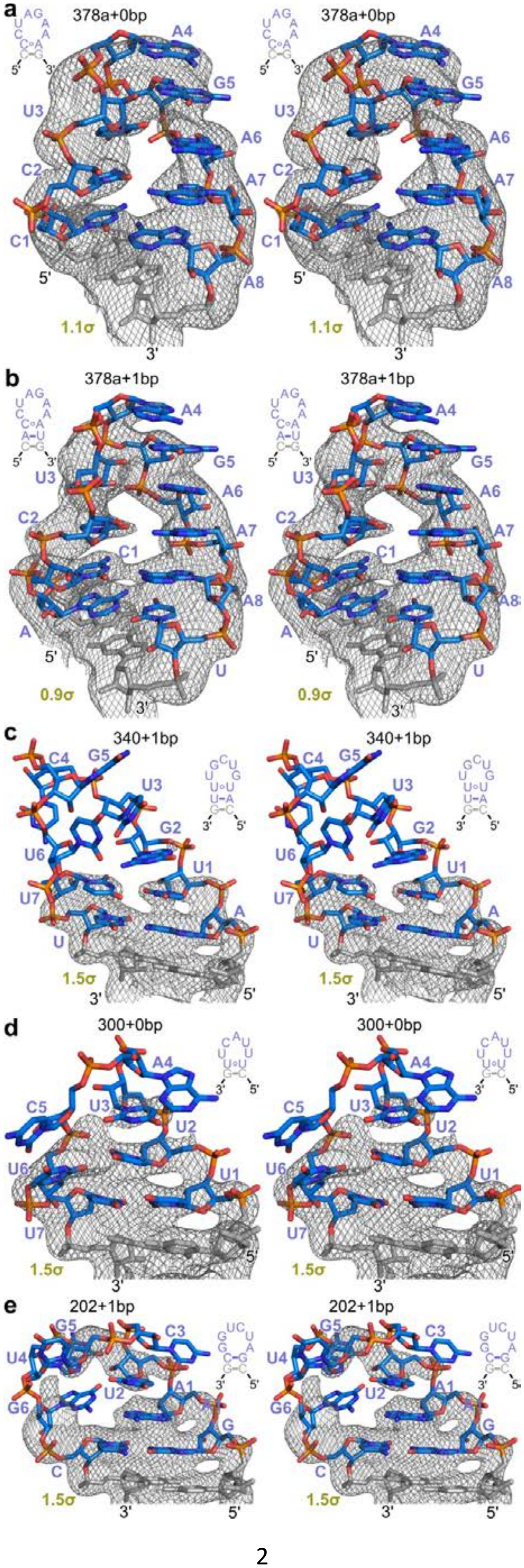

a
378a+0bp
378a+0bp
A4
G5
U3
A6
C2
A7
C1
A8
5'
3'
1.1σ
b
378a+1bp
378a+1bp
A4
G5
U3
A6
C2
A7
C1
A8
A
U
5'
3'
0.9σ
c
340+1bp
340+1bp
C4
G5
U3
G2
U6
U1
U7
A
U
1.5σ
3'
5'
d
300+0bp
300+0bp
A4
C5
U3
U2
U6
U1
U7
1.5σ
3'
5'
e
202+1bp
202+1bp
G5
C3
U4
U2
A1
G6
G
C
1.5σ
3'
5'

**Supplementary Fig. 1** $\sigma_A$-weighted 2Fo-Fc simulated annealing omit maps of pri-miRNA apical junctions and loops 6-8 nt in length. Color scheme is the same as Fig. 2. See Methods for details on map calculation.

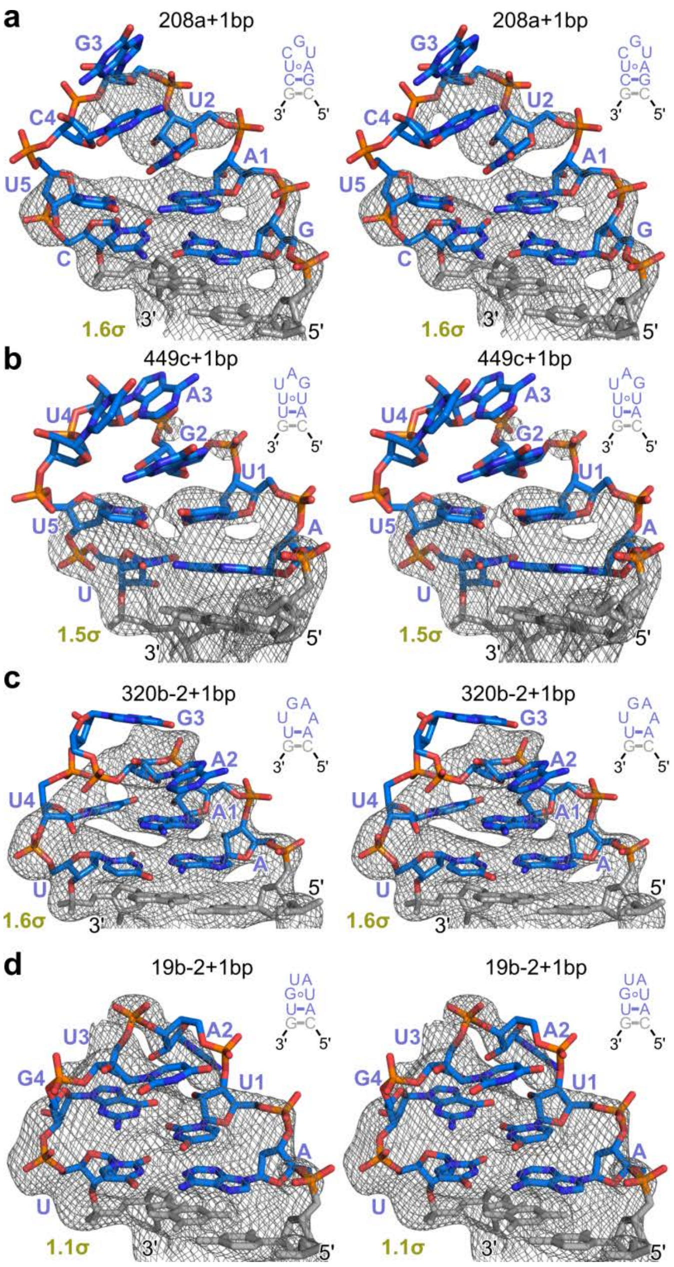


**Supplementary Fig. 2** $\sigma_A$-weighted 2Fo-Fc simulated annealing omit maps of pri-miRNA apical junctions and loops 4-5 nt in length. Color scheme is the same as Fig. 2. See Methods for details on map calculation.

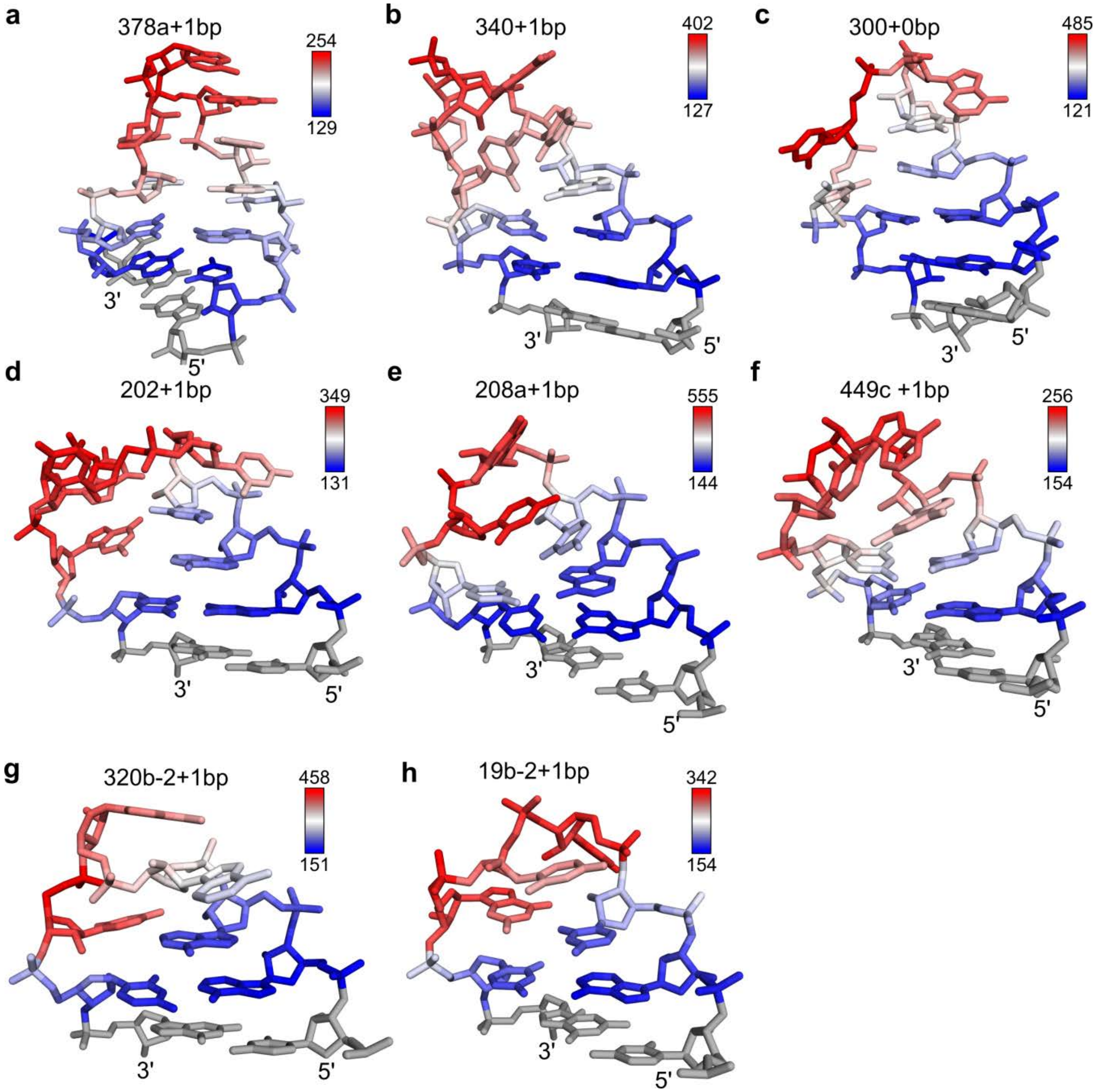


**Supplementary Fig. 3** miR-precursor apical loop structures colored with atomic displacement parameters. The insets show the ADP ranges plotted.

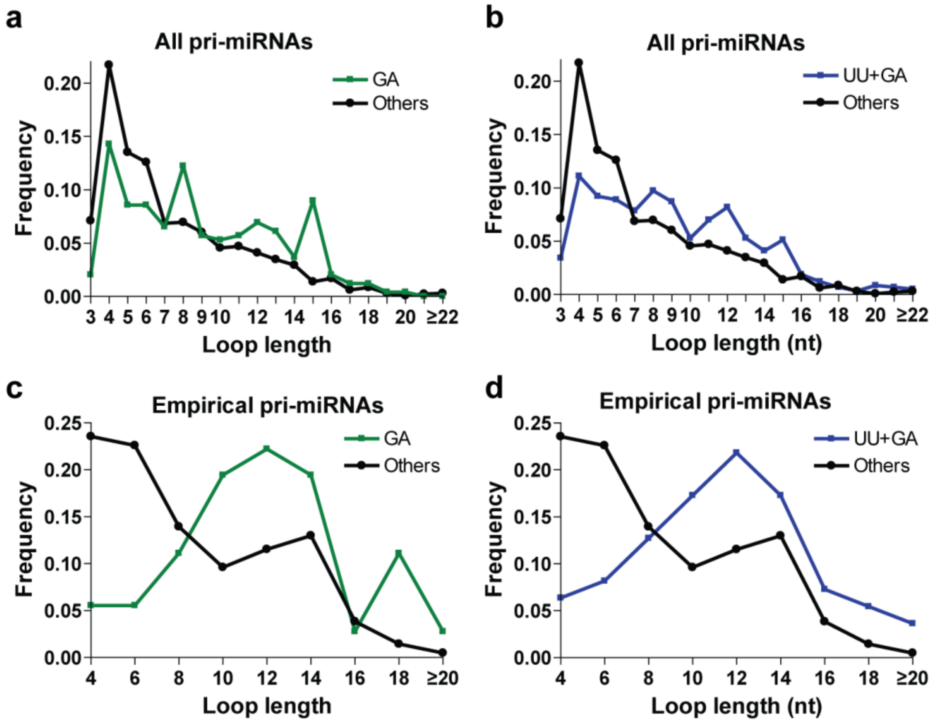


**Supplementary Fig. 4** Loop length distributions of pri-miRNAs containing (analysis extended from Fig. 4g,h). "UU+GA" combines the pri-miRNAs containing UU and GA. "Others" represents the pri-miRNAs that contain neither UU nor GA pair at the apical junction. Either all pri-miRNAs (**a**, **b**) or the empirical set (**c**, **d**) are used.

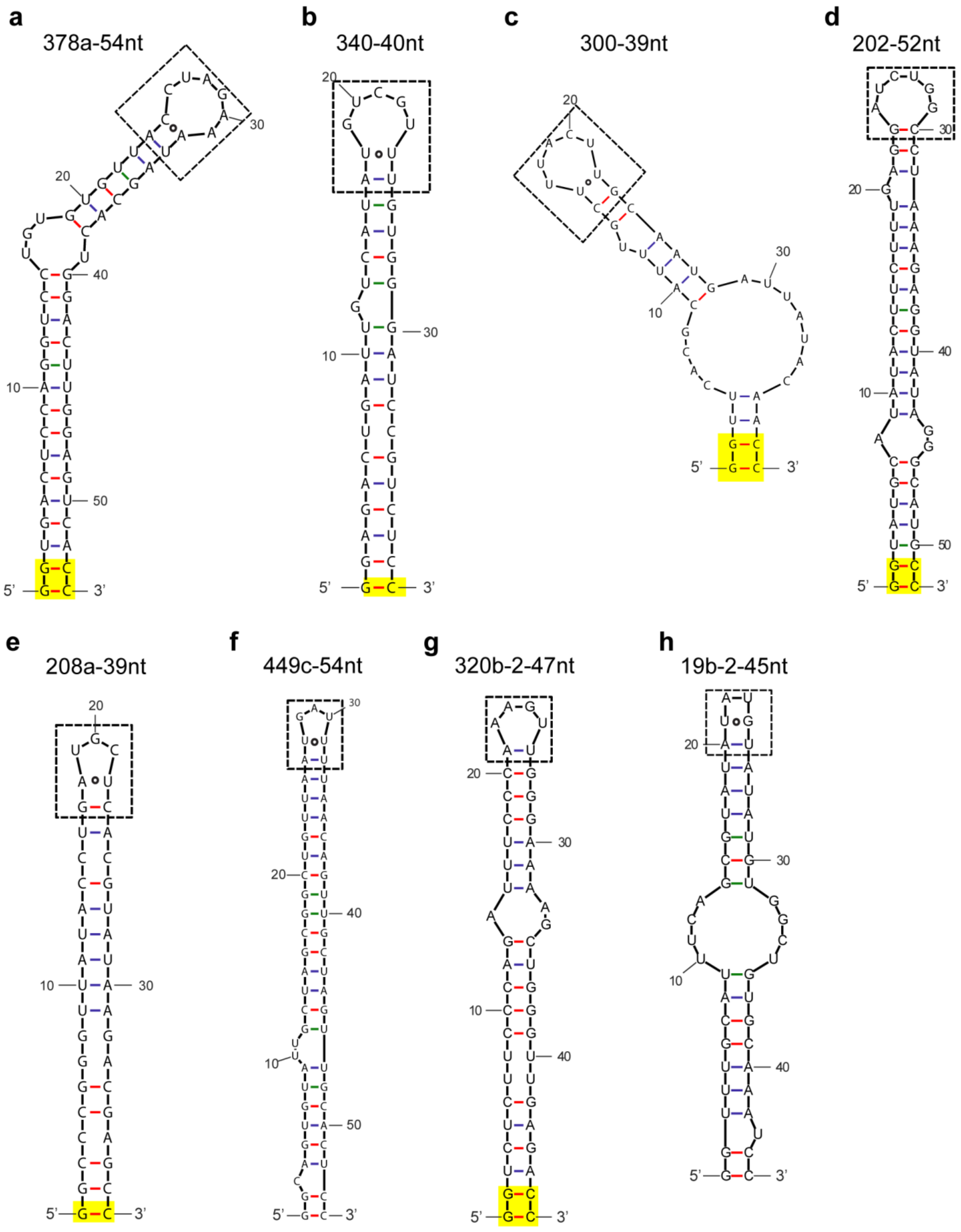


**Supplementary Fig. 5** RNA constructs used in Rhed-binding assays. Additional G-C pairs added to the base of the stem to enhance transcription are highlighted in yellow. The boxes show the apical loops and terminal stem base pairs for which crystal structures have been determined.

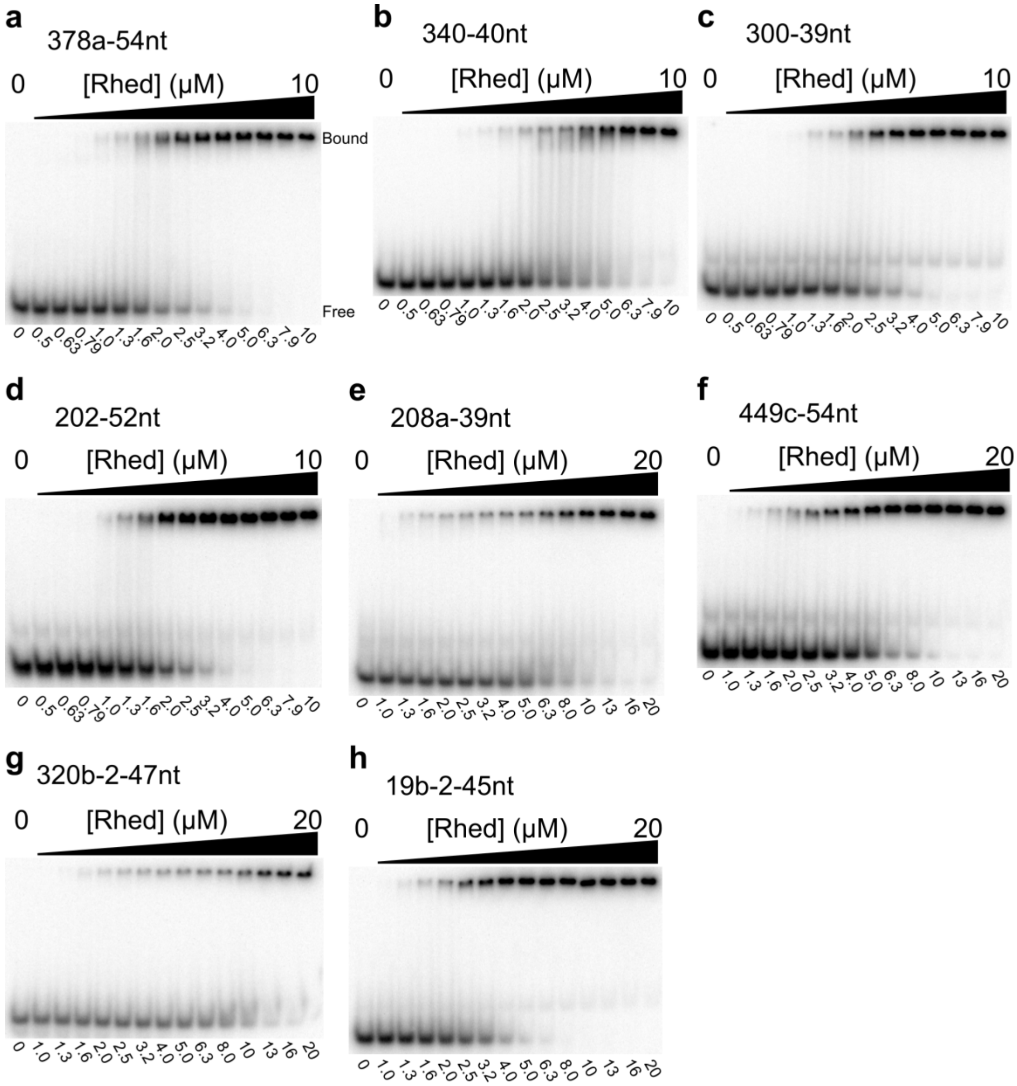


**Supplementary Fig. 6** Example electrophoresis mobility shift assays for measuring binding affinities of pri-miRNA fragments to Rhed. The free RNA and protein-bound species are labeled in **a**. Rhed dimer concentrations (in μM) used in the binding reactions are shown below the gels. The Rhed-bound bands are in the gel and are not aggregates stuck in wells.

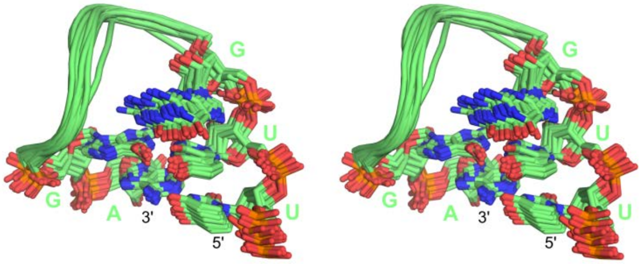


**Supplementary Fig. 7** Stereo diagram of NMR ensemble for pre-miR-20b, showing the U-G pair at the apical junction and stacking of the neighboring 5' G residue [1].

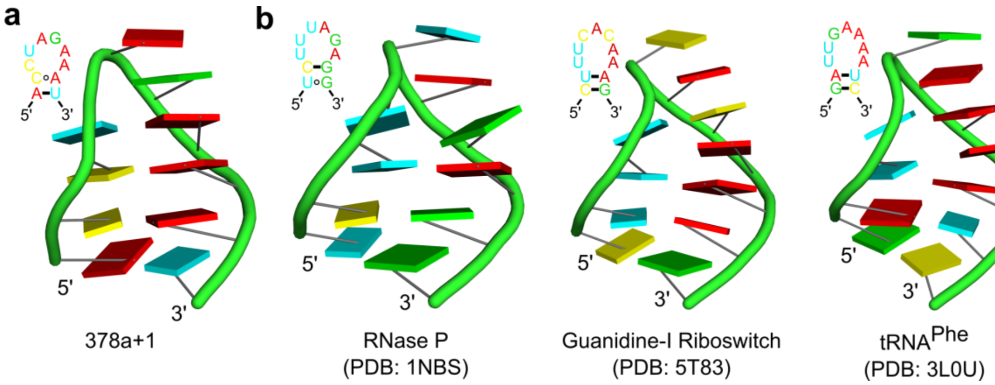


**Supplementary Fig. 8** Comparison of pri-miRNA terminal loop structures to similar RNA folds found in the PDB. Cartoon representation of (**a**) the 8-nt loop of pri-miR-378a (378a+1, left), (**b**) similar loops from the structures of RNase P [2], guanidine-I riboswitch [3], and tRNA$^{Phe}$ [4].
